# Single cell analyses reveal phosphate availability as critical factor for nutrition of *Salmonella enterica* within mammalian host cells

**DOI:** 10.1101/2020.10.23.351551

**Authors:** Jennifer Röder, Michael Hensel

## Abstract

*Salmonella enterica* serovar Typhimurium (STM) is an invasive, facultative intracellular pathogen that resides in a specialized membrane-bound compartment termed *Salmonella*-containing vacuole (SCV). Essential for survival and proliferation in the SCV is *Salmonella* pathogenicity island II (SPI2) that encodes a type III secretion system (T3SS). The SPI2-T3SS and the effector translocation maintain SCV integrity and formation of specific tubular membrane compartments, called *Salmonella*-induced filaments (SIFs). The SCV/SIF continuum allows STM to bypass nutritional restriction in the intracellular environment by acquiring nutrients from the host cell. Phosphate is one of the most abundant elements in living organisms and in STM, inorganic phosphate (P_i_) homeostasis is mediated by the two-component regulatory system PhoBR, resulting in expression of the high affinity phosphate transporter *pstSCAB-phoU.* Using fluorescent protein reporters, we investigate P_i_ availability for STM at single cell level over time within the intracellular habitats of different host cells. We observed that the *pstSCAB-phoU* encoded phosphate uptake system is essential for intracellular replication of STM because there is a P_i_ ion concentration less 10 μM within the SCV. Additionally, the demand and consumption of P_i_ correlates with intracellular proliferation of STM and we identify a dependency of SPI2 activity and P_i_ starvation.

## Introduction

Microbial pathogens have to adapt to the specific nutritional landscape provided by the host in order to survive and to proliferate. For intracellular bacteria, nutritional conditions within host cells may vary depending on the specific subcellular niche inhabited by the pathogen, the antimicrobial activity of the host cell, and the pathogen burden. Given the current antibiotics crisis due to increasing resistance and lack of novel antimicrobial compounds, bacterial nutrition within the host may provide new targets for non-antibiotic strategies to combat infectious diseases. Understanding bacteria nutrition within the host and/or host cells is prerequisite for such approaches, and we set out to investigate the nutritional requirements of *Salmonella enterica*, a food-borne pathogen causing diseases ranging from gastroenteritis to systemic typhoid fever.

*S. enterica* serovar Typhimurium (STM) is an invasive, facultative intracellular pathogen that resides in a specialized membrane-bound compartment termed *Salmonella*-containing vacuole (SCV) (Haraga, Ohlson, & Miller, 2008). Essential for survival and proliferation in the SCV is a type III secretion system (T3SS) encoded by *Salmonella* pathogenicity island 2 (SPI2) (Hensel et al., 1998). The SPI2-T3SS and the translocated effector proteins maintain the SCV integrity and mediate formation of specific tubular membrane compartments, called *Salmonella*-induced filaments (SIFs) (Brumell, Tang, Mills, & Finlay, 2001; Drecktrah, Knodler, Howe, & Steele-Mortimer, 2007; Figueira & Holden, 2012; Rajashekar, Liebl, Seitz, & Hensel, 2008). Formation of an extensive SIF network by STM was suggested to provide bacterial nutrition and enabling proliferation (Liss et al., 2017). Mutant strains defective in SPI2-T3SS are attenuated in systemic virulence and show reduced intracellular replication (Hensel et al., 1995; Hensel et al., 1998). An important SPI2-T3SS effector protein is SifA, as lack of this effector leads to defects in remodeling of the host cell endosomal system and maintenance of intact SCV (Beuzon et al., 2000; Stein, Leung, Zwick, Garcia-del Portillo, & Finlay, 1996). Deletion of *sifA* attenuates virulence, replication in macrophages (Beuzon et al., 2000; Brumell, Rosenberger, Gotto, Marcus, & Finlay, 2001), and results in hyper-replication of STM in the nutrient-rich cytosol of epithelial cells (Knodler, 2015).

Intracellular STM show a remarkable heterogeneity regarding proliferation and response to environmental cues (Bumann & Cunrath, 2017; Helaine et al., 2014). Subpopulations are present in SCV or cytosol, are actively proliferating or dormant (Helaine et al., 2014; Knodler, 2015). For analyses of the intracellular environment of STM and responses to intracellular cues, this heterogeneity restricts the application of population-wide approaches such as transcriptomics or proteomics. Such analyses on the level of single intracellular bacteria are still pending. However, bacteria can be used as sensitive tools to interrogate the response to specific environmental cues, and fluorescent protein reporters in combination with flow cytometry (FC) allows efficient single cell analyses of large and heterogenous populations. We set out to systematically analyze the availability of nutrients for intracellular STM in mammalian host cells on the single cell level. Here, we focus on the role of phosphate for intracellular STM. Phosphorus is one of the most abundant elements in living organisms, found in membrane lipids, nucleic acids, proteins or carbohydrates, and involved in many enzymatic reactions. In bacteria, inorganic phosphate (P_i_) homeostasis is mediated by the two-component regulatory system PhoBR. The membrane-bound sensor PhoR reacts to low P_i_ and activates PhoB, a transcriptional regulator that promotes the expression of many genes involved in P_i_ transport (Hsieh & Wanner, 2010). This includes transcription of the *pstSCAB-phoU* operon encoding a high affinity phosphate transporter. PstS is a periplasmic protein that binds P_i_ with high affinity. PstC, PstA and PstB form an ABC transporter with PstC and PstA being inner membrane channel proteins and PstB an ATP-dependent permease component. PhoU is essential for the suppression of the Pho regulon under a high phosphate condition (Wanner, 1996). The exact mechanism of action of PhoU is not yet understood (Gardner, Johns, Tanner, & McCleary, 2014). Several studies link the P_i_ homeostasis with virulence of bacteria as well as for STM (Lamarche, Wanner, Crepin, & Harel, 2008).

Expression of SPI2-T3SS genes is activated within the SCV of host cells, and the proper spatiotemporal control ensures activity of virulence factors in the required phase of host-pathogen interaction. The intracellular habitat of STM is a complex environment with a multiplicity of stressors and nutritional limitations, thus specific stimuli are difficult to dissect. In contrast, systematic analyses of expression under defined *in vitro* conditions identified a slightly acidic pH, Mg^2+^ limitation, and P_i_ starvation as stimuli inducing SPI2 gene expression (Deiwick & Hensel, 1999). The SPI2-T3SS-dependent endosomal remodeling and induction of a complex SCV/SIF continuum allows intracellular STM to relieve nutritional limitations (Liss et al., 2017). Accordingly, we anticipate a link between nutritional limitation in the intracellular environment, control of SPI2 gene expression, and SPI2-T3SS-mediated manipulation of the host cell.

Phosphate uptake and the regulation of P_i_ homeostasis by *pstSCAB-phoU* were suggested to be essential for the pathogenicity of STM. Various studies indicated upregulation during infection (Garcia-del Portillo, Foster, Maguire, & Finlay, 1992; Hautefort et al., 2008). Thus, we investigated P_i_ availability for STM at a single cell level over time within distinct intracellular habitats of different host cells. Therefore, we generated reporters and measured the presence and concentration of P_i_ during infection of host cells by STM.

## Results

### The *pstSCAB phoU*-encoded phosphate uptake system is required for intracellular proliferation of STM

We investigated the relevance of P_i_ uptake for the intracellular lifestyle of STM. Various genes involved in regulation of P_i_ homeostasis were deleted and intracellular proliferation of the resulting mutant strains in comparison to STM WT were determined using competitive index (CI) assays (**Figure 1**). The SPI2-T3SS-deficient strain Δ*ssaV* served as negative control for intracellular replication (Hensel et al., 1998) and showed a low CI of 0.1, both in HeLa cells and RAW264.7 macrophages. The *pstSCAB* operon encodes for an ABC transporter, mediating the phosphate uptake (Lamarche et al., 2008). Deletion of this transporter resulted in a CI of 0.4 in both host cell types. This high-affinity phosphate transporter is regulated by the *phoBR* two-component system (Lamarche et al., 2008). Deletion of *phoB* or *phoR* attenuated intracellular replication. In HeLa cells, CI of 0.2 or 0.3 were determined for Δ*phoB or* Δ*phoR* strains, respectively, and within RAW264.7 macrophages as host cells, CI of 0.5 or 0.6 for Δ*phoB* or Δ*phoR* strains, respectively, were determined.

**Figure 1.**
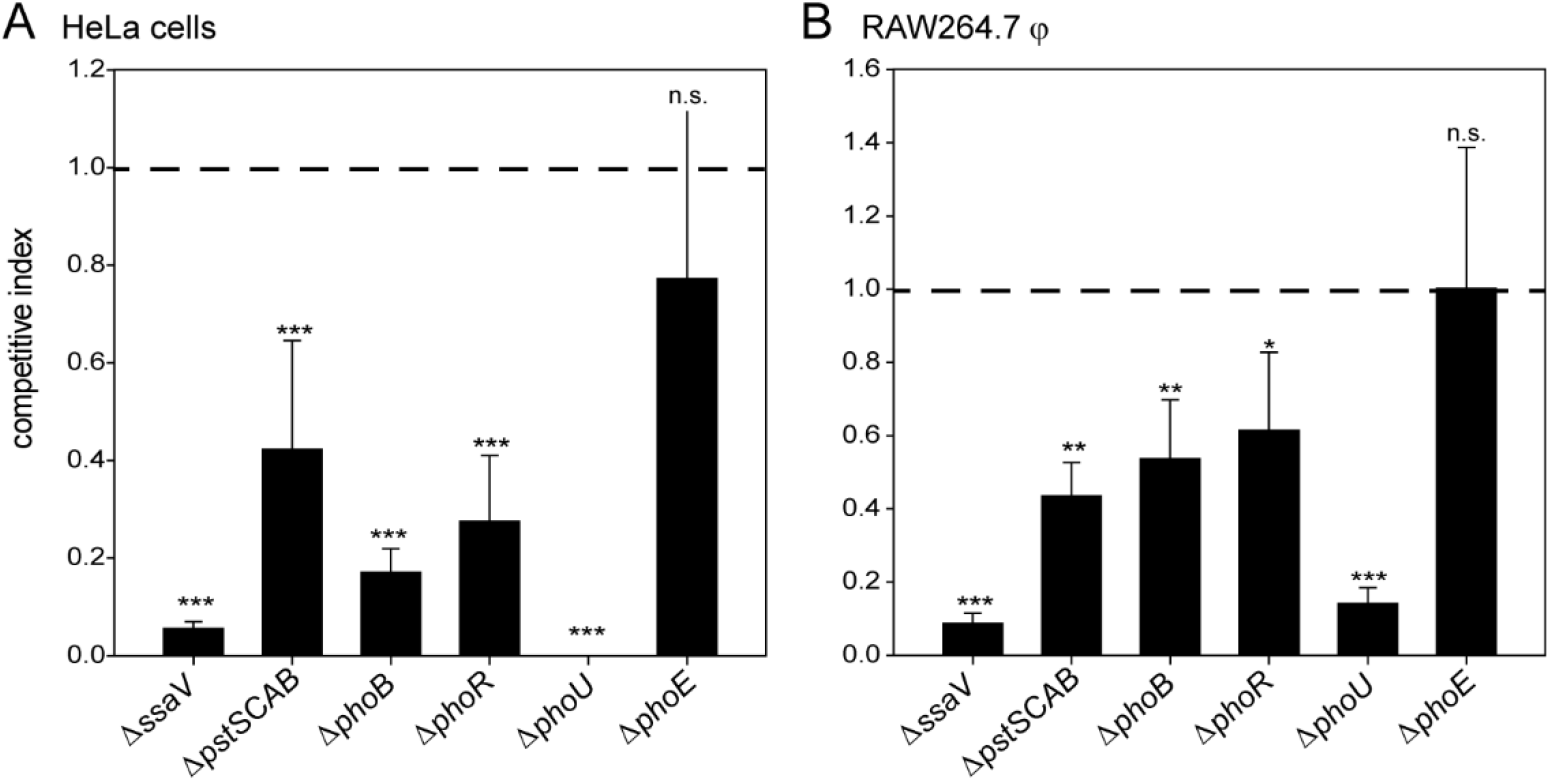
Role of phosphate transport systems for intracellular proliferation of STM. Competitive index (CI) assays for proliferation of STM strains in HeLa cells (A), or RAW264.7 macrophages (B). Cells were co-infected with STM WT in combination with mutant strains defective in *ssaV*, *pstSCAB*, *phoB*, *phoR*, *phoU*, or *phoE* at a total multiplicity of infection (MOI) of 1. Intracellular proliferation was determined as the ratio of CFU at 16 h p.i. to CFU at 1 h p.i. and CI of replication of WT versus mutant strain was calculated. A CI of 1.0 (dashed line) indicates identical intracellular proliferation of both strains, a CI lower than 1.0 indicates attenuated proliferation of the mutant strain. Shown are mean values and standard deviations of three biological replicates, each consisting of three technical replicates. Statistical analyses were performed by one-way ANOVA in comparison to STM WT and are expressed as: n.s., not significant; *, p <0.05; **, p < 0.01; ***, p < 0.001.

A particularly interesting phenotype was observed by deletion of *phoU.* While in RAW264.7 macrophages phagocytosis was not affected, replication was impaired with a CI of 0.1. A Δ*phoU* strain showed reduced growth in LB medium and formed smaller colonies compared to STM WT. PhoU plays a key role in phosphate homeostasis by repressing the *pho* regulon at high phosphate levels (Gardner et al., 2014). SPI1-T3SS-mediated invasion of HeLa cells was below the detection limit, probably due to the reduced growth rate. This resulted in a highly reduced CI for Δ*phoU* vs. WT in HeLa cells. In contrast, lack of PhoE, an outer membrane protein for uptake of inorganic phosphates (Spierings, Elders, van Lith, Hofstra, & Tommassen, 1992), did not affect intracellular replication as CI of 0.7 and 1 in HeLa and RAW264.7, respectively, were determined. Therefore, we conclude that proper P_i_ homeostasis plays a central role for intracellular replication and virulence of STM.

### Generation and validation of reporter strains for measuring inorganic phosphate levels

Phosphate uptake by the transporter PstSCAB, as well as regulation of this system by PhoBR, proved to be of critical importance for intracellular proliferation. This prompted us to investigate availability of P_i_ for intracellular STM in more detail. Intracellular STM diversify into various subpopulations, i.e. bacteria being located vacuolar or cytosolic, proliferating slowly or rapidly, or forming non-replicating persisters. This heterogeneity demands analyses on level of single cell of intracellular STM. In order to determine the intracellular P_i_ concentration more precisely, we generated a dual fluorescence reporter with constitutive expression of DsRed, and sfGFP under control of the promoter of *pstS*. PstS is a periplasmic protein that binds P_i_ with high affinity and is known to be upregulated intracellularly in P_i_-poor environments (Hautefort et al., 2008).

STM was subcultured for 3.5 h or 24 h in media with various concentrations of P_i_, and expression of P_*pstS*_∷sfGFP was determined by FC. Induction of P_*pstS*_∷sfGFP was detected after growth in media containing P_i_ concentrations of 1 mM or lower, and sfGFP intensity increased as P_i_ concentration decreased (**Figure 2**AB). Comparison of P_*pstS*_∷sfGFP intensities between cells of 3.5 h and 24 h subcultures revealed higher intensity in late cultures (**Figure 2**C). This would be in line with the consumption of the P_i_ pool during STM growth resulting in P_i_ limitation, and/or accumulation of the reporter. To precisely determine the P_i_ concentration critical for induction of phosphate reporter P_*pstS*_∷sfGFP, we performed a phosphate shock experiment (**Figure 2**DE). A logarithmic culture was used to inoculate media with various P_i_ concentrations and incubation was continued for 1 h. This regime minimized the change in P_i_ levels due to consumption by bacterial growth, and we detected induction of the phosphate reporter at concentrations below 100 μM. Decreasing P_i_ concentrations correlated with increasing sfGFP intensities, and sfGFP intensity of 10^4^ relative fluorescence intensity (RFI) indicated a concentration of less than 10 μM P_i_ (**Figure 2**F). The effect of mutations in phosphate transporters or regulatory systems on the expression of the phosphate reporter was analyzed (Figure S 1). In PCN medium containing 25 mM P_i_, here referred to as PCN (25), no induction of P_*pstS*_∷sfGFP was detected. However, a Δ*phoU* strain demonstrated strong induction of P_*pstS*_∷sfGFP in PCN (0.4), as well as in PCN (25) despite sufficient phosphate availability. Only moderate growth was observed for this mutant strain, indicating missing repression of *pstS* in the Δ*phoU* background. While P_*pstS*_∷sfGFP in STM WT showed increased intensity from PCN (25) to PCN (0.4), Δ*pstSACB*, Δ*phoB* and Δ*phoR* strains did not respond to low phosphate concentrations. Therefore, no regulation of the high-affinity P_i_ uptake system was possible in these mutant strains.

**Figure 2.**
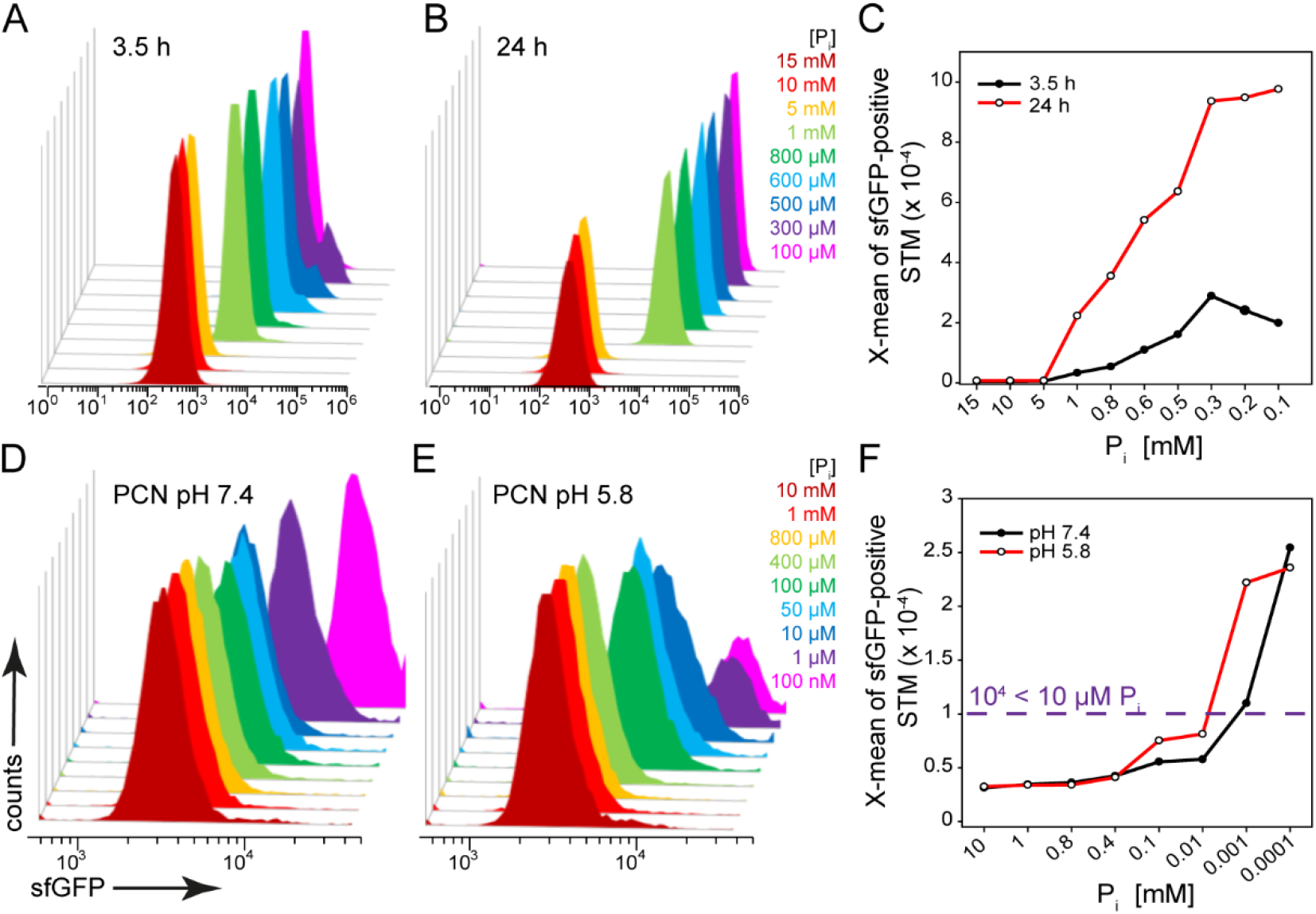
A dual fluorescence reporter for measuring the presence of phosphate. STM harboring p5007 for constitutive expression of DsRed, and sfGFP under control of P_*pstS*_ were cultured in PCN minimal medium with various amounts of P_i_. Induction of P_*pstS*_∷sfGFP *in vitro* was determined by flow cytometry (FC). A, B, C) STM WT [p5007] was grown overnight (o/n) in PCN containing 25 mM P_i_ (PCN (25)), pH 7.4 and then subcultured 1:31 in PCN, pH 7.4 with various concentrations of P_i_ as indicated. Samples were collected after 3.5 h (A, C black line) or 24 h (B, C red line) and X-means of sfGFP intensity of P_*pstS*_-induced bacteria was determined by FC (C). D, E, F) STM WT [p5007] was grown o/n in PCN (25) pH 7.4, diluted 1:100 in PCN (1), pH 7.4 or PCN (1), pH 5.8, and subcultured to OD_600_ 0.5-0.7. These subcultures were used to inoculate PCN media adjusted to pH 7.4 (D, F black line) or 5.8 (E, F, red line) with various concentrations of P_i_ as indicated, and culture was continued for 1 h, and X-means of sfGFP intensities were determined (F). sfGFP intensities above 10^4^ RFI indicate P_i_ concentrations lower than 10 μM. sfGFP intensities of P_*pstS*_-positive bacteria of a representative experiment are shown.

### Heterogeneity of the intracellular bacterial populations

After validation of the phosphate reporter, we deployed the system in infection assays of HeLa cells and RAW264.7 macrophages. Due to the different sfGFP intensities, we detected heterogeneous STM populations in HeLa cells and RAW264.7 macrophages. After infection of HeLa cells, three subpopulations were detected at 8 h post-infection (p.i.) (**Figure 3**ABC). The classification of P_i_ availability was based on the intensities determined by *in vitro* experiments. A very small population P1 was non-induced, a large population P2 (ca. 90%) indicated P_i_ concentrations above 10 μM, and a small population P3 (4%) corresponded to concentrations below 10 μM P_i_. At 16 h p.i., P3 of STM WT (34%) and Δ*ssaV* strains (18.5%) increased, while the Δ*sifA* strain remained predominantly in P2 (88%). Since a *sifA* mutant strain has access to cytosolic components due to compromised SCV integrity (Beuzon et al., 2000), we assumed a higher P_i_ concentration in the host cell cytosol. Therefore, we deduced for HeLa cells P_i_ concentrations lower than 10 μM in the SCV, and higher than 10 μM in the cytosol. In RAW264.7 macrophages, conditions were slightly different (**Figure 3**DEF). At 8 h p.i. we distinguished between induced (55% of STM WT and 52% of Δ*ssaV*) and non-induced populations, but all induced STM reported presence of less than 10 μM P_i_. While the Δ*ssaV* strain showed 70% P_*pstS*_-positive bacteria at 16 h p.i., the STM WT showed 92% of P_*pstS*_-positive bacteria at 16 h p.i., including 25% of cells with lower DsRed and sfGFP intensity, indicative for concentrations of more than 10 μM P_i_.

**Figure 3.**
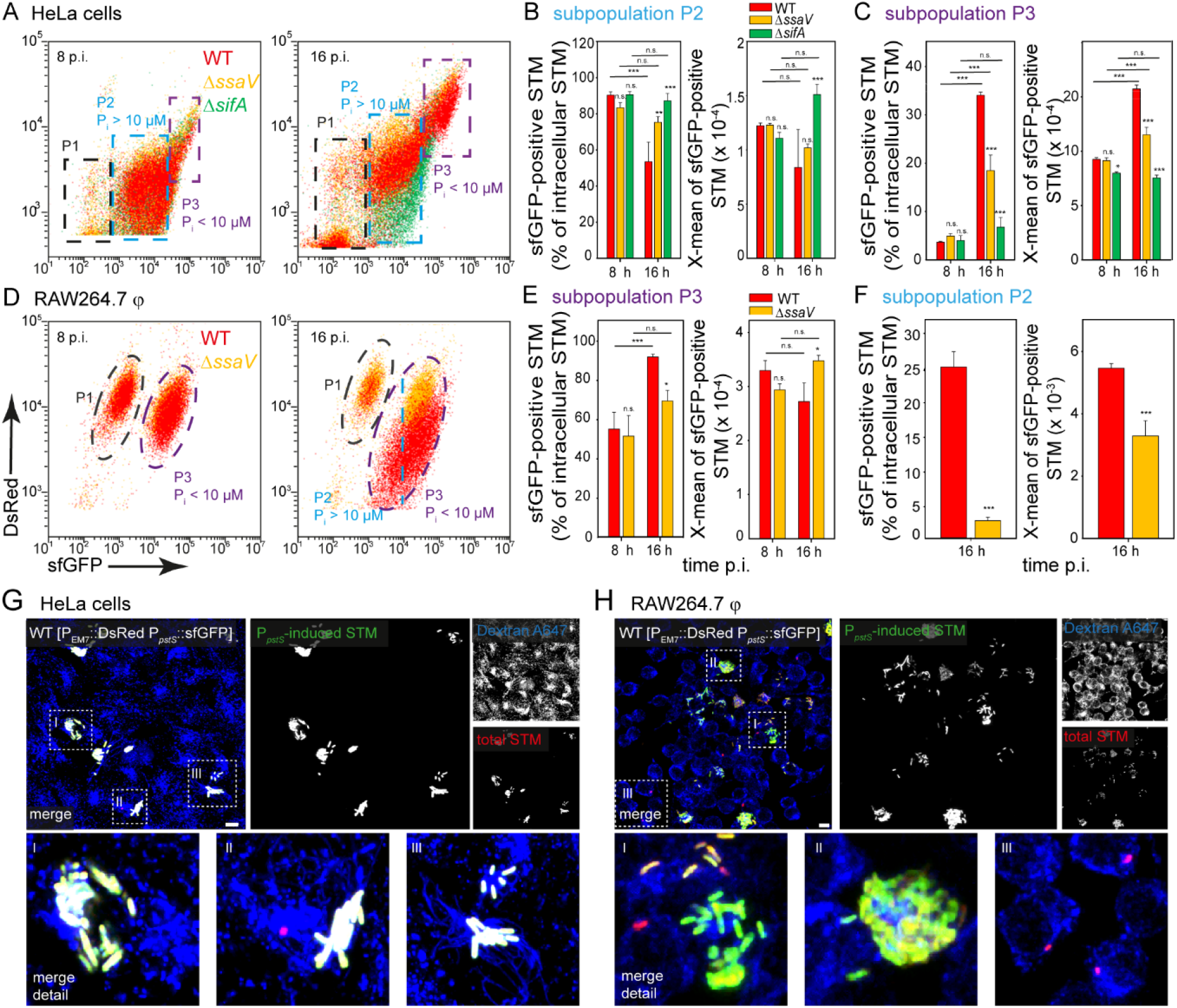
Determination of phosphate availability for intracellular STM in HeLa cells or RAW264.7 macrophages. HeLa cells (A-C) or RAW264.7 macrophages (D-F) were infected at MOI 5 with STM WT (red), Δ*ssaV* (orange) or Δ*sifA* (green) strains as indicated, each containing the phosphate reporter p5007. A, B) Host cells were lysed 8 h or 16 h p.i., released STM were fixed and subjected to FC to quantify sfGFP intensities of P_*pstS*_-positive STM. Three subpopulations of intracellular STM were distinguished based on P_*pstS*_∷sfGFP intensity: P1, P_*pstS*_-negative; P2, P_*pstS*_-positive at P_i_ concentration > 10 μM; and P3, P_*pstS*_-positive at P_i_ concentration < 10 μM. Representative quantification of population size and X-means of sfGFP intensities of subpopulations P2 (B, F) and P3 (C, E) for STM WT, Δ*ssaV* and Δ*sifA* strains in HeLa cells (B, C), and STM WT and Δ*ssaV* strains in RAW264.7 macrophages (E, F) at 8 h and 16 h p.i. Mean values and standard deviations of P_*pstS*_-positive bacterial subpopulations from triplicates of a representative experiment are shown. Statistical analyses were performed by one-way ANOVA for mutant strains compared to STM WT, or between time points, and are expressed as: n.s., not significant; *, p < 0.05; **, p < 0.01; ***, p < 0.001. HeLa cells (G) or RAW264.7 macrophages (H) were infected with STM WT [p5007] and pulse-chased with dextran-AlexaFluor 647 (blue) for labelling of the endosomal system. Live cell imaging was performed 16 h p.i., and overview images show heterogeneous sfGFP (green) intensities of STM. Representative infected cells indicate red and green fluorescence signal for STM. Sections in the dashed box are shown magnified below. Scale bars, 10 μm.

In overview images, STM WT indicated quite evenly distributed sfGFP intensities in HeLa cells (**Figure 3**G). However, in RAW264.7 macrophages the heterogeneity of DsRed and sfGFP intensities was more visible (**Figure 3**H). The Δ*ssaV* strain (Figure S 3) also showed a relatively uniform distribution of sfGFP intensity in HeLa cells. In RAW264.7 macrophages, we distinguished between non-induced and induced bacteria, whereas the sfGFP intensity did not exhibit major differences.

We tested another phosphate reporter using the promoter of *apeE*, which codes for an outer membrane esterase. Expression of *apeE* was induced by phosphate limitation, and required the *phoBR* phosphate regulatory system (Conlin, Tan, Hu, & Segar, 2001). This reporter led to comparable results. An *in vitro* induction was measured at concentrations below 1 mM in a 3.5 h subculture and the sfGFP intensity increased with decreasing P_i_ values (Figure S 2A). The same subpopulations within HeLa cells (Figure S 2BCD) and RAW264.7 macrophages (Figure S 2EF) were observed, but generally the sfGFP intensity was lower.

Next, we analyzed induction of the phosphate reporter over the time course of infection. By comparing the reporter induction of STM WT and Δ*ssaV* strains in HeLa cells, we determined similar induction up to 10 h p.i., as percentage of sfGFP-positive STM and intensity of single bacteria increased over time (**Figure 4**AB). P_*pstS*_-positive bacteria increased from 5% at 2 h p.i. to 85% for STM WT, and to 62% for the Δ*ssaV* strain at 10 h p.i. At later time points, the Δ*ssaV* strain decreased to 34% P_*pstS*_-induced cells, while STM WT continued to increase to 89% (**Figure 4**CD). This likely reflects the higher replication and metabolic activity of STM WT compared to STM Δ*ssaV* (Liss et al., 2017).

**Figure 4.**
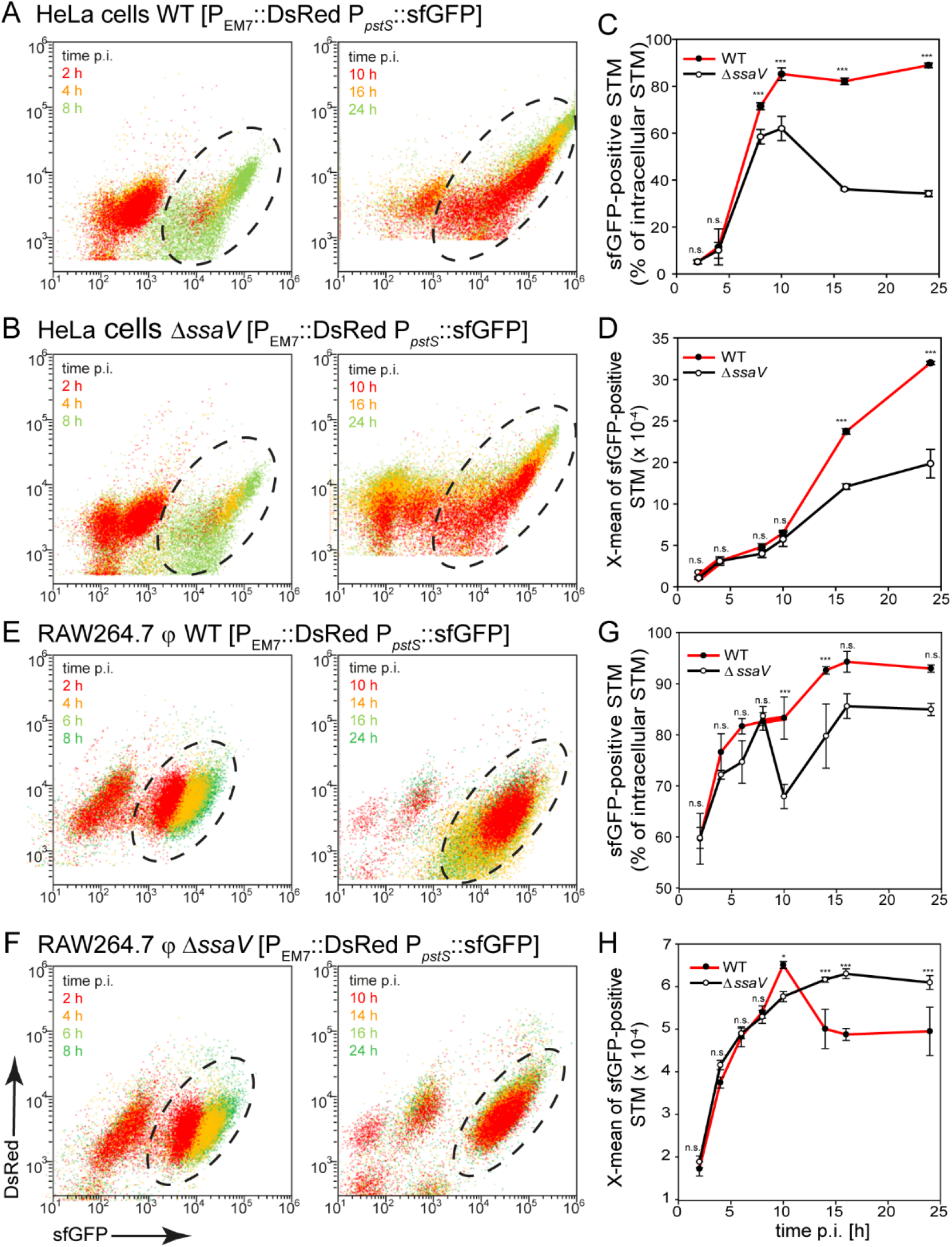
Kinetics of phosphate availability for STM in HeLa cells and RAW264.7 macrophages. HeLa cells (A-D) or RAW264.7 macrophages (E-H) were infected at MOI 5 with STM WT (A, E) or Δ*ssaV* (B, F) strains each harboring p5007. Host cells were lysed at various time points p.i. as indicated, released bacteria were fixed and subjected to FC to quantify the population of P_*pstS*_-positive STM (C, G), and X-means of sfGFP intensities (D, H) for STM WT (red lines) and Δ*ssaV* (black lines) strains in HeLa cells or RAW264.7 macrophages. Mean values and standard deviations from triplicates of a representative experiment are shown. Statistical analyses are indicated as for **Figure 3**.

A steady increase of P_*pstS*_-positive cells was observed for STM isolated from RAW264.7 macrophages over time (**Figure 4**EF). With an increase from 60% at 2 h p.i. to 93% and 85% for STM WT and Δ*ssaV* strains, respectively, at 24 h p.i. The Δ*ssaV* strain exhibited slight fluctuations, but over time between 60-85% of the bacteria were induced. In contrast, STM WT showed a decline in sfGFP intensity at 10 h and later p.i. (**Figure 4**GH). Therefore, in HeLa cells P3 (< 10 μM P_i_) became larger, and P2 (> 10 μM P_i_) smaller over time, because of increasing sfGFP intensities. In RAW264.7 macrophages, a severe P_i_ deficiency was already detected in the early stages, and again replicating STM WT differ from non-replicating Δ*ssaV* strain after 10 h p.i.

Due to the different populations observed for STM WT and Δ*sifA* strains in HeLa cells, we already assumed that access to cytosolic components led to increased phosphate availability. The use of a dual reporter with P_*uhpT*_∷DsRed and P_*pstS*_∷sfGFP provided information on phosphate concentration over time in distinct habitats in HeLa cells (**Figure 5**A, Figure S 5A). P_*uhpT*_ induction indicates cytosolic presence of the bacteria (Röder & Hensel, 2020). After invasion, most bacteria are still in a vacuole and therefore P_*uhpT*_-negative. However, the population of P*uhpT-*positive bacteria increased in the early intracellular phase, i.e. by replication in the cytosol. At late time points (≥10 h p.i.) change in intracellular population occurred, with an increased vacuolar population, and declined of the cytosolic population. For STM WT, the P_*uhpT*_-negative population increased from 54% at 10 h p.i. to 81% at 24 h p.i. The sfGFP intensity increased, indicating decreased availability of P_i_ in both habitats (**Figure 5**BC). Similar results were obtained with a reporter using the correlation between P_*ssaG*_∷DsRed and P_*pstS*_∷sfGFP (**Figure 5**D, Figure S 5B). The *ssaG* gene is located in SPI2, encodes the needle subunit of the T3SS, and was used to analyze expression of genes for the SPI2-T3SS (Lim, Kim, Choi, Lee, & Ryu, 2006). SPI2 induction is known for SCV-bound bacteria, but not for cytosolic bacteria (Knodler et al., 2010). Accordingly, vacuolar replication also started 10 h p.i., as the P_*ssaG*_-positive population increased from 39% to 75% at 24 h p.i., but unlike the P_*uhpT*_-positive population the intensity of the P_*ssaG*_-negative population did not increase (**Figure 5**EF). This may be attributed to the continuously rupture of the SCV by STM (Röder & Hensel, 2020), i.e. STM escaped from a P_i_-restricted SCV into the cytosol of host cells, and thus erroneously enhanced the overall sfGFP intensity. Equally, cytosolic bacteria (P_*ssaG*_-negative) are unlikely to become vacuolar and exhibited continuously high sfGFP intensity. In RAW264.7 macrophages, however, all bacteria were P_*ssaG*_-positive (vacuolar), and only few P_*uhpT*_-positive bacteria (cytosolic) were detected (Figure S 4). Therefore, STM mainly resides inside the SCV of RAW264.7 macrophages.

**Figure 5.**
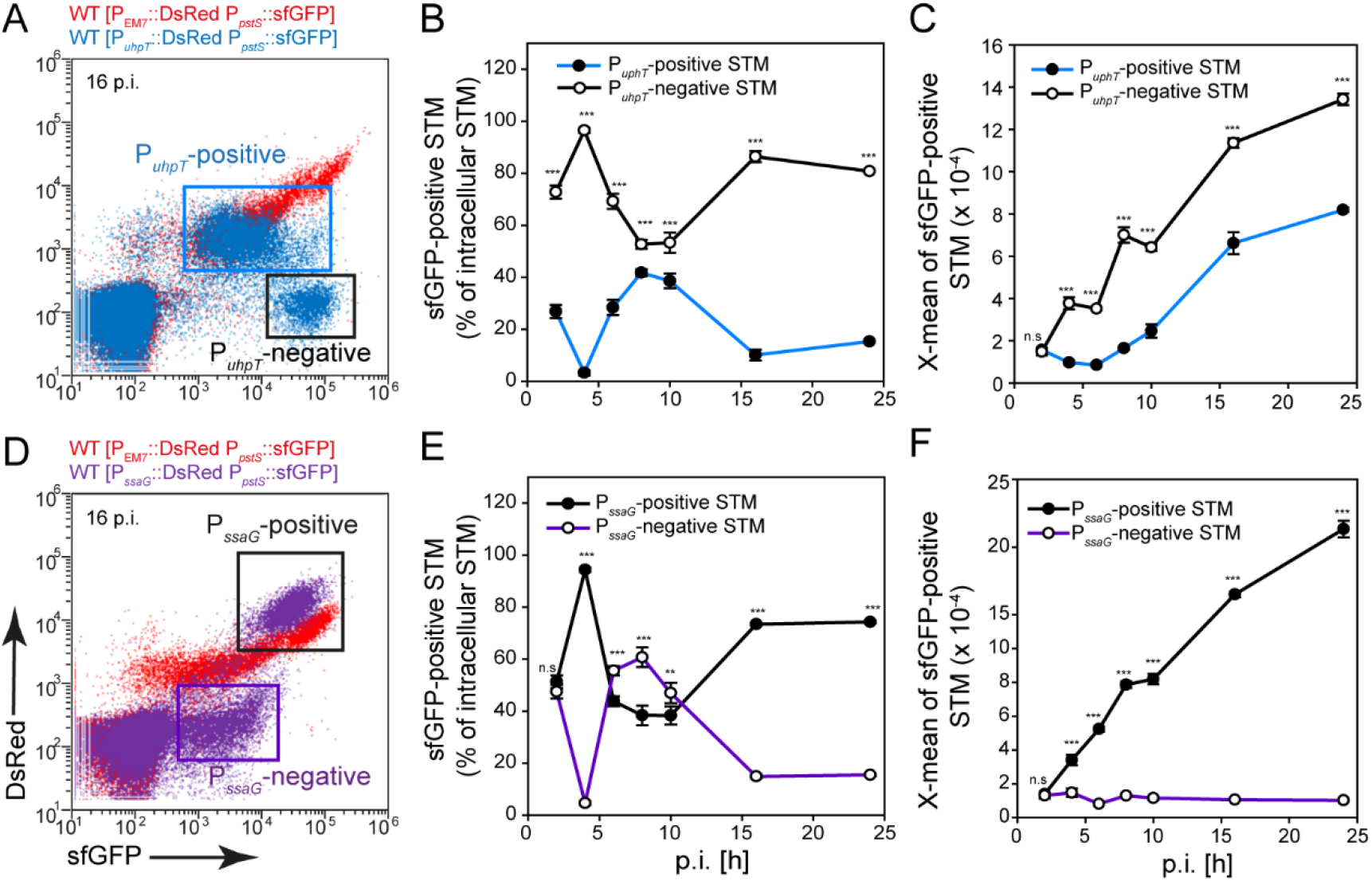
Higher phosphate availability for STM in host cell cytosol. HeLa cells were infected at MOI 5 with STM WT harboring p5193 with DsRed under control of P_*uhpT*_ and P_*pstS*_∷sfGFP (blue), or p5457 for expression of DsRed under control of P*ssaG,* and P_*pstS*_∷sfGFP (violet). For comparison and gating of populations, HeLa cells were infected with STM WT harboring p5007 for constitutive expression of DsRed and P_*pstS*_∷sfGFP (red). Host cells were lysed at various points in time p.i. as indicated, released STM were fixed and subjected to FC for quantification of P_*pstS*_-positive bacteria. Induction of P_*uhpT*_ or P_*ssaG*_ was determined. Data for STM WT [p5007] and WT [p5193] (A), or STM WT [p5007] and WT [p5457] (D) of a representative experiment are shown. The sizes (B, E) and sfGFP intensities (C, F) of the cytosolic STM population (P_*uhpT*_∷sfGFP-positive, blue lines, P_*ssaG*_∷sfGFP-negative, violet lines), and SCV-bound STM population (P_*uhpT*_∷sfGFP-negative, P_*ssaG*_∷sfGFP-positive, black lines) were quantified and mean values and standard deviations for represent data from triplicates are shown. Statistical analyses are indicated as for **Figure 3**.

We also analyzed the phosphate reporter in activated RAW264.7 macrophages and primary human macrophages, which provided a more restrictive environment for bacteria resulting in an attenuated replication (Lathrop et al., 2018; Rosenberger & Finlay, 2002). If RAW264.7 macrophages activated with interferon-γ (IFN–γ), were infected, we obtained mixed populations for STM WT or Δ*ssaV* strains (**Figure 6**A). The induced bacterial population was smaller (70% and 95% in activated and non-activated RAW264.7, respectively). However, the intensity was higher, suggesting a lower P_i_ concentration (**Figure 6**B).

**Figure 6.**
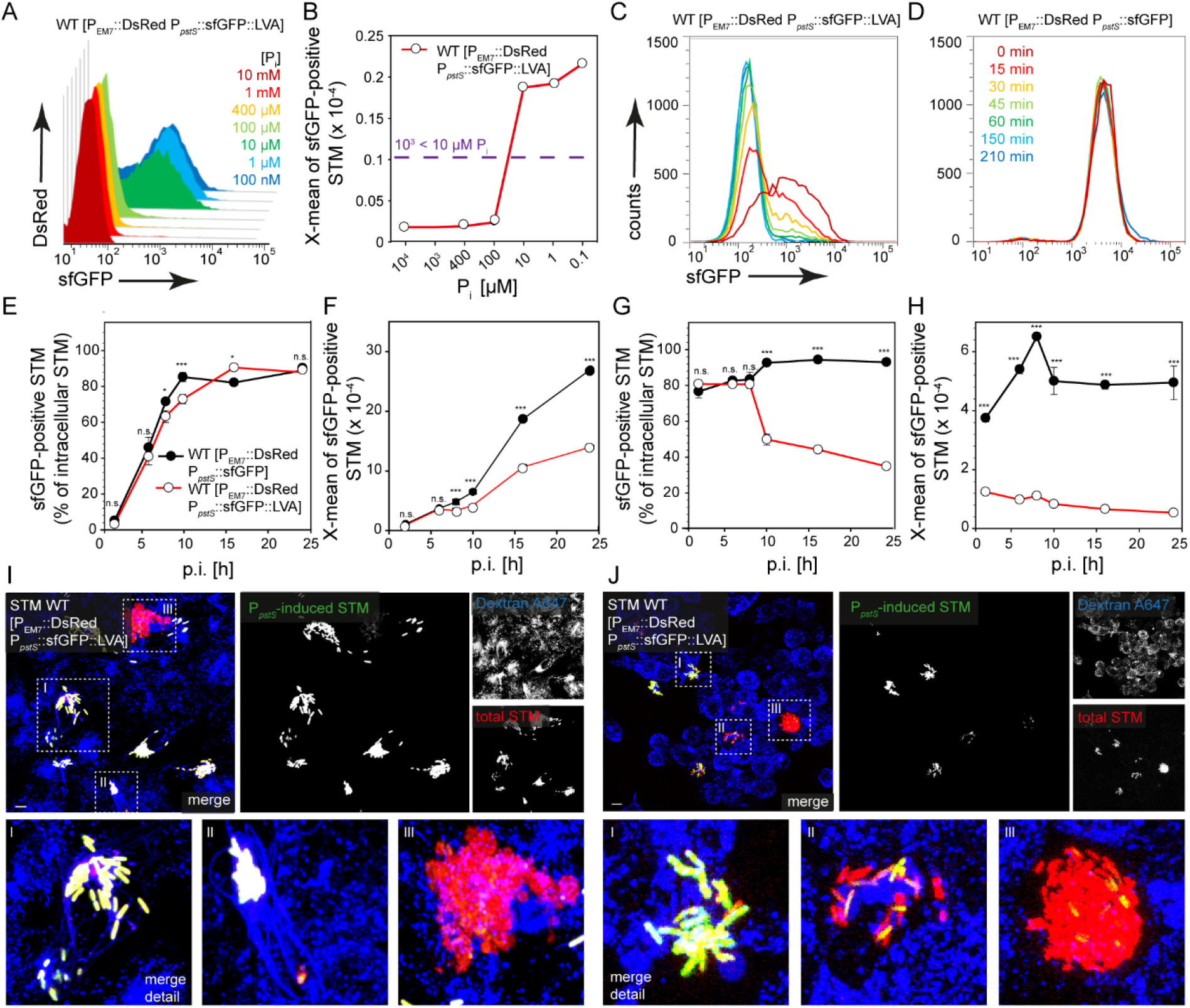
Phosphate limitation for STM in IFN-γ-activated RAW264.7 macrophages and monocyte-derived human macrophages. A, B) RAW264.7 macrophages were cultured for 24 h in medium without (blue) or with (red) 5 ng x ml^−1^ IFN-γ, and subsequently infected with STM WT [p5007] at MOI 5. C, D) Human peripheral blood macrophages (red, orange) or RAW264.7 macrophages (light green, dark green) were infected with STM WT [p5007] at MOI 25. Host cells were lysed as indicated at 8 h p.i. or 16 h p.i., released STM were fixed and subjected to FC to quantify P_*pstS*_-positive STM and X-means of sfGFP intensities for P_*pstS*_-positive STM (B, D). Data of representative experiments are shown, (A) for STM WT [p5007] in resting RAW264.7 (red) and activated RAW264.7 (blue), (C) for STM WT [p5007] in RAW264.7 (light green, dark green). or human macrophages (red, orange) Mean values and standard deviations from triplicates of a representative experiment are shown. Statistical analyses are indicated as for **Figure 3**.

In primary macrophages isolated from human peripheral blood we determined an even smaller induced population (32% and 26% at 8 h and 16 h p.i., respectively), but it is known that proliferation for STM in these cells is highly restricted. Since only about 30% of the bacteria were induced, the rest of the population may be dead, or have entered persister state (**Figure 6**CD). Although the population was much more widely distributed in RAW264.7 macrophages, a similar average sfGFP intensity was measured at 16 h p.i., whereas at 8 h p.i. the sfGFP intensity was much lower in human macrophages. However, in activated RAW264.7 macrophages, as well as in primary human macrophages, induction and thus phosphate limitation was detected.

### Reporters with destabilized sfGFP allow to measure rapid changes in phosphate availability

The sfGFP has a rather long half-life (Pedelacq, Cabantous, Tran, Terwilliger, & Waldo, 2006) that limits analyses of STM responses to changing environments. To modify the reporter system for analyses of rapid changes in phosphate concentration, the LVA tag was fused to sfGFP resulting in increased degradation (Andersen et al., 1998). Analysis of induction *in vitro* in a phosphate shock experiment showed that slightly delayed induction occurred and signals were measured at a concentration of 10 μM P_i_, but not at 100 μM (**Figure 7**A). At a concentration of 10 μM P_i_ an intensity of 10^3^ RFI was determined for sfGFP-LVA, compared to 10^4^ RFI for the parental reporter, probably as consequence of the continuous degradation of sfGFP (**Figure 7**B). Therefore, we compared the stability of sfGFP and sfGFP-LVA by growing subcultures in PCN (0.01) media for 2 h. Then, protein synthesis was stopped by chloramphenicol, and incubation continued. The sfGFP intensities of the cultures were determined at various time points after block of synthesis (**Figure 7**CD), and we observed that the LVA tag caused a continuous decline of sfGFP intensity and no sfGFP signals were detectable after 60 min. For normal sfGFP, a constant signal was still detectable after 210 min.

**Figure 7.**
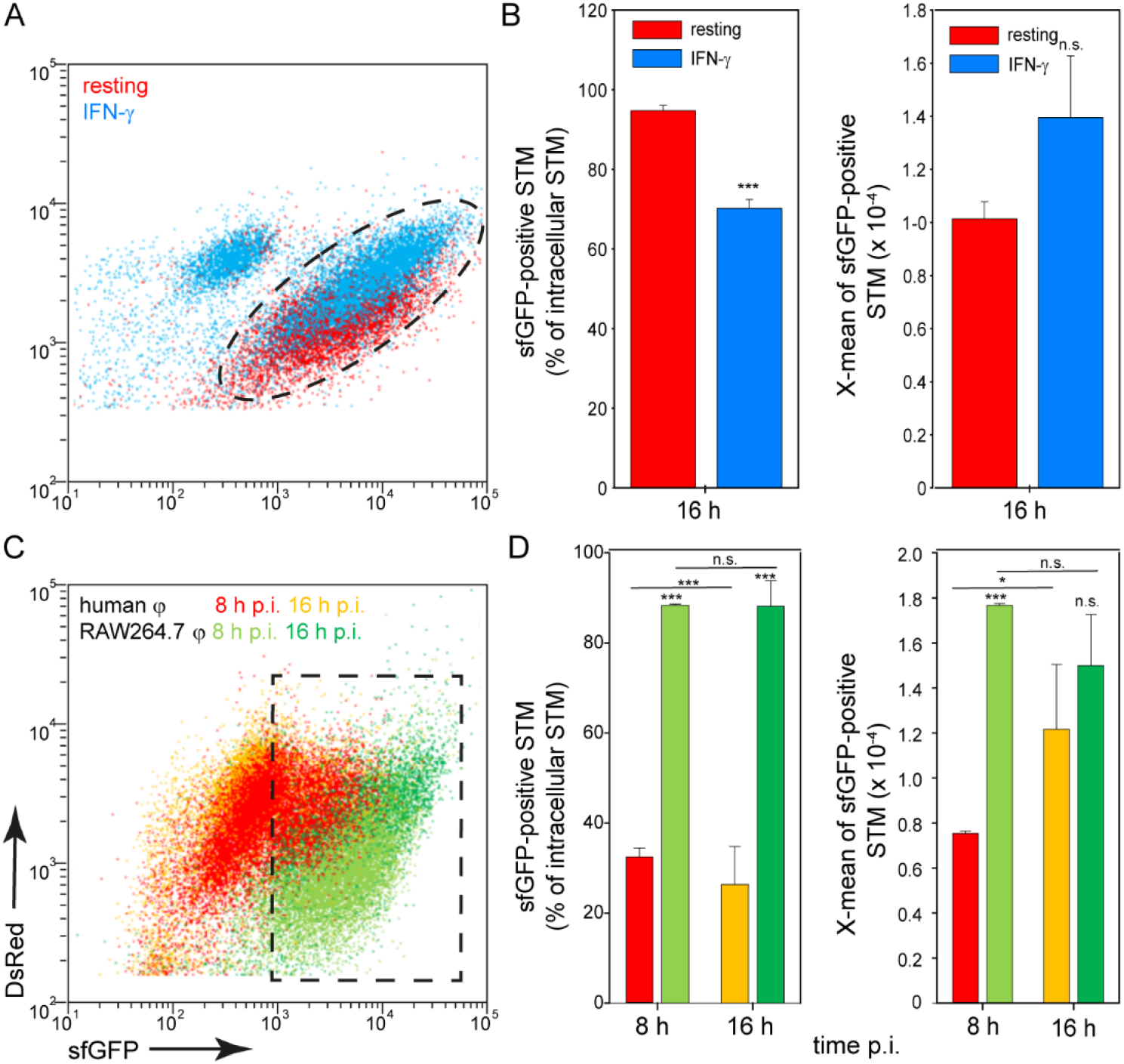
Analyses of dynamics in phosphate availability using a reporter with destabilized sfGFP. STM harbored p5440 for constitutive expression of DsRed, and sfGFP∷LVA under control of P_*pstS*_. A) WT [p5440] was grown o/n in PCN (25) pH 7.4, diluted 1:100 in PCN (1), pH 5.8, and subcultured to OD_600_ 0.5-0.7. This culture was used to inoculate PCN media with various concentrations of P_i_ as indicated, and culture was continued for 1 h. B) The sfGFP intensities of P_*pstS*_-induced STM was determined by FC. Phosphate concentrations lower than 10 μM P_i_ corresponded to sfGFP intensities higher than 10^3^ RFI. sfGFP intensities of P_*pstS*_-positive bacteria of a representative experiment are shown. C, D) The stability of sfGFP and sfGFP∷LVA was determined. WT [p5007] or WT [p5440] were diluted 1:31 in PCN (0.01), pH 5.8 and subcultured for 2 h. Chloramphenicol was added to 200 ng x ml^−1^ final concentration to inhibit bacterial protein synthesis. The culture was further incubated, and sfGFP intensities were determined at various culture times as indicated. HeLa cells (E, F) or RAW264.7 macrophages (G, H) were infected with STM WT [p5007] or WT [p5440] at MOI 5. Host cells were lysed at various time points p.i., released STM were fixed and subjected to FC to quantify P_*pstS*_-positive bacteria and sfGFP intensities. Mean values and standard deviations of triplicates of a representative experiment are shown. Statistical analyses are indicated as for **Figure 3**. HeLa cells (I) or RAW264.7 macrophages (J) were infected with STM WT [p5440] and pulse-chased with dextran-AlexaFluor 647 for labelling of the endosomal system. Live cell imaging was performed 16 h p.i. Representative infected cells showed red and green fluorescence signals for STM. Sections in the dashed box are shown magnified below. Scale bars, 10 μm.

We used this reporter in STM infection of HeLa cells and observed hardly any difference between P_*pstS*_∷sfGFP and P_*pstS*_∷sfGFP-LVA (**Figure 7**EF). The percentages of induced bacteria were almost identical, while sfGFP intensity was lower if LVA was present. Nevertheless, the intensity of the destabilized sfGFP constantly increased, thus confirming a decrease in P_i_ concentration over time of intracellular proliferation. In RAW264.7 macrophages, sfGFP intensity dropped after 10 h p.i. both for sfGFP without and with LVA tag (**Figure 7**GH). With LVA labeling, however, not only the change in the intensity was detected, but also the proportion of P_*pstS*_-positive bacteria highly decreased. The overview images showed that especially in heavily loaded HeLa cells and RAW264.7 macrophages, no further sfGFP signals were visible for LVA-tagged version (**Figure 7**IJ).

### Intracellular proliferation limits phosphate availability for STM

We found that STM WT generally exhibited stronger P_*pstS*_∷sfGFP induction in HeLa cells than in RAW264.7 macrophages. Furthermore, the percentage of P_*pstS*_-induced bacteria was lower in activated RAW264.7 macrophages and in human macrophages compared to resting host cells, or in STM Δ*ssaV* strain compared to STM WT. This might be explained by different composition of compartments in distinct cells types, or by different levels of STM proliferation affecting intracellular P_i_ levels due to P_i_ consumption. To investigate the influence of replication on P_i_ availability we infected HeLa cells or RAW264.7 macrophages and added Cotrimoxazole to inhibit intracellular replication. In HeLa cells we observed no changes at 8 h p.i., but significantly lower sfGFP intensity at 16 h p.i. indicating higher P_i_ availability (**Figure 8**ABC). Quantification of CFU of intracellular STM indicated lower bacterial counts in Cotrimoxazole-treated cells at 8 h and 16 h p.i. (**Figure 8**D). No significant increase in CFU was observed, showing that STM remained viable but restricted in proliferation. In RAW264.7 macrophages, both the percentage and the intensity were lower at 16 h p.i. due to inhibition of bacterial replication (**Figure 8**EFG). Cotrimoxazole-treatment also reduced CFU of STM in macrophages, but proliferation between 8 h and 16 h p.i. was not fully ablated (**Figure 8**H). However, Cotrimoxazole-inhibited STM in macrophages showed a lower frequency of sfGFP-positive cells at 16 h p.i., with sfGFP intensities as low as at 8 h p.i. We conclude that inhibition of intracellular proliferation relieves the limitation of P_i_ due to decreased consumption of the P_i_ pool available for intracellular STM.

**Figure 8.**
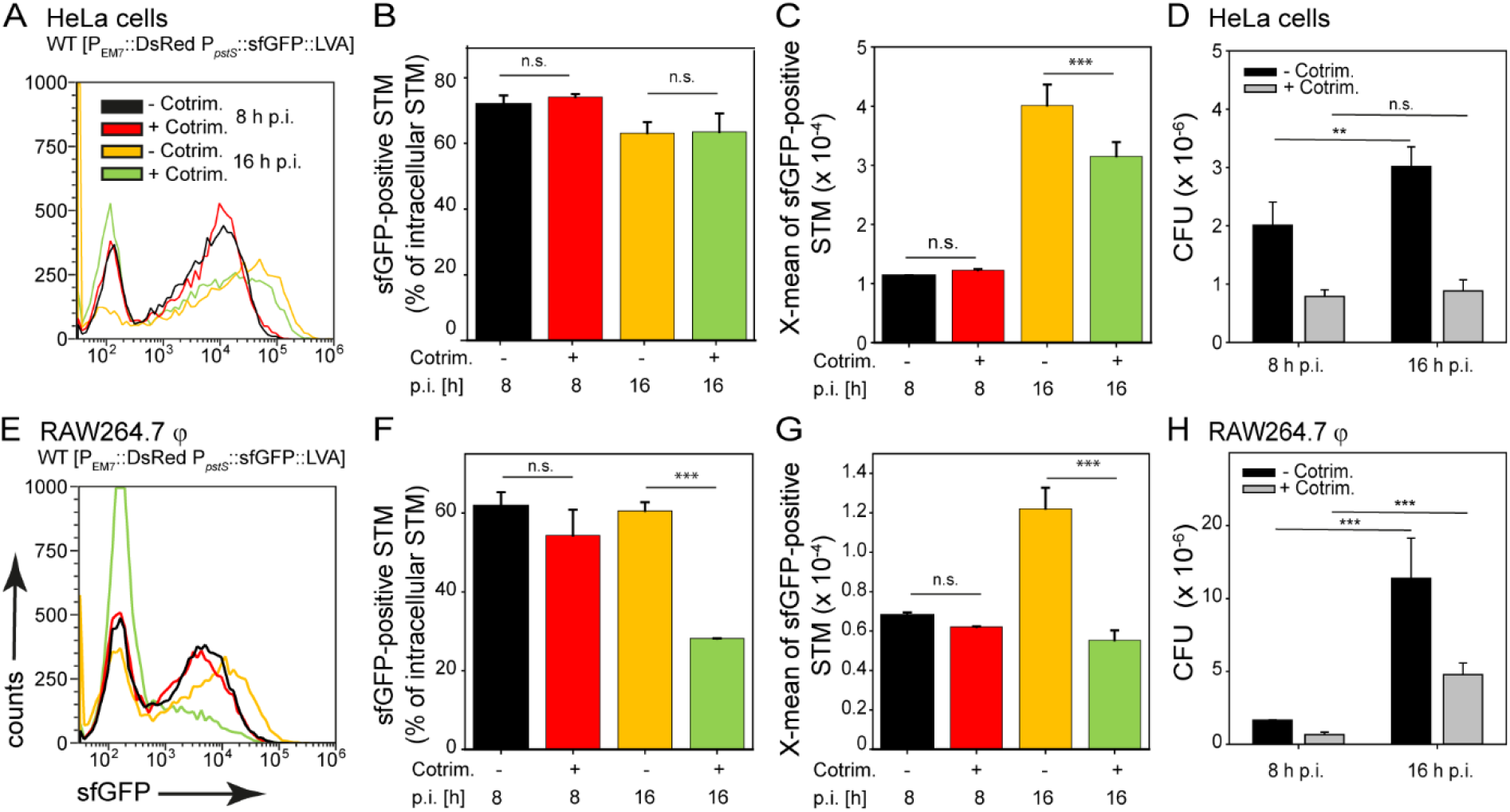
Inhibition of bacterial proliferation ablates intracellular phosphate limitation. HeLa cells (A-D) or RAW264.7 macrophages (E-H) were infected with STM WT [p5440] at MOI 5. If indicated, Cotrimoxazole was added at 1 h p.i. to 100 ng x ml^−1^ final concentration to infected cells in order to arrest STM intracellular proliferation. Host cells were lysed 8 h p.i. (black, red) or 16 h p.i. (orange, green), released STM were fixed and subjected to FC for quantification of P_*pstS*_-positive STM (B, F), and X-means for sfGFP intensities of P_*pstS*_-positive STM (C, G). Representative data are given for STM WT in nontreated or Cotrimoxazole-treated HeLa cells (A) or RAW264.7 macrophages (E). Mean values and standard deviations of triplicates of a representative experiment are shown. D, H) Host cells were lysed 8 h or 16 h p.i., released STM were plated onto agar plate to determine CFU of intracellular STM in nontreated (black bars) and in Cotrimoxazole-treated (grey bars) HeLa cells (D), or RAW264.7 macrophages (H). Statistical analyses are indicated as for **Figure 3**.

Finally, we manipulated phosphate availability in host cells and determined the effects on intracellular STM. For this, i) additional phosphate was added to the cell culture medium to increase P_i_ uptake (Candeal, Caldas, Guillen, Levi, & Sorribas, 2014), ii) acidification of the SCV was inhibited by bafilomycin (Rathman, Sjaastad, & Falkow, 1996), or iii) the proton gradient was uncoupled by protonophore CCCP (Candeal et al., 2014). We added P_i_ to the host cells 24 h before infection, so that a higher P_i_ level was already present in the host cells. When we infected HeLa cells with STM WT, fewer bacteria showed induction of P_*pstS*_∷sfGFP (**Figure 9**ABC). In infected RAW264.7 macrophages additional P_i_ had no effect on P_*pstS*_∷sfGFP expression by intracellular STM (**Figure 9**DEF). In contrast, the addition of bafilomycin and thus blocking the acidification of the SCV increased sfGFP intensities of STM in RAW264.7 macrophages but not for STM in HeLa cells. Uncoupling by CCCP resulted in slightly higher sfGFP intensity of STM in HeLa cells, but no changes were detected in RAW264.7 macrophages. In conclusion, the higher external P_i_ levels affect P_i_ availability of STM in HeLa cell, while neutralization of the pH of SCV in RAW264.7 macrophages increased P_i_ limitation, which may be a consequence of reduced transport of P_i_ into the SCV lumen, and/or higher P_i_ consumption by increased STM proliferation in a more permissive compartment.

**Figure 9.**
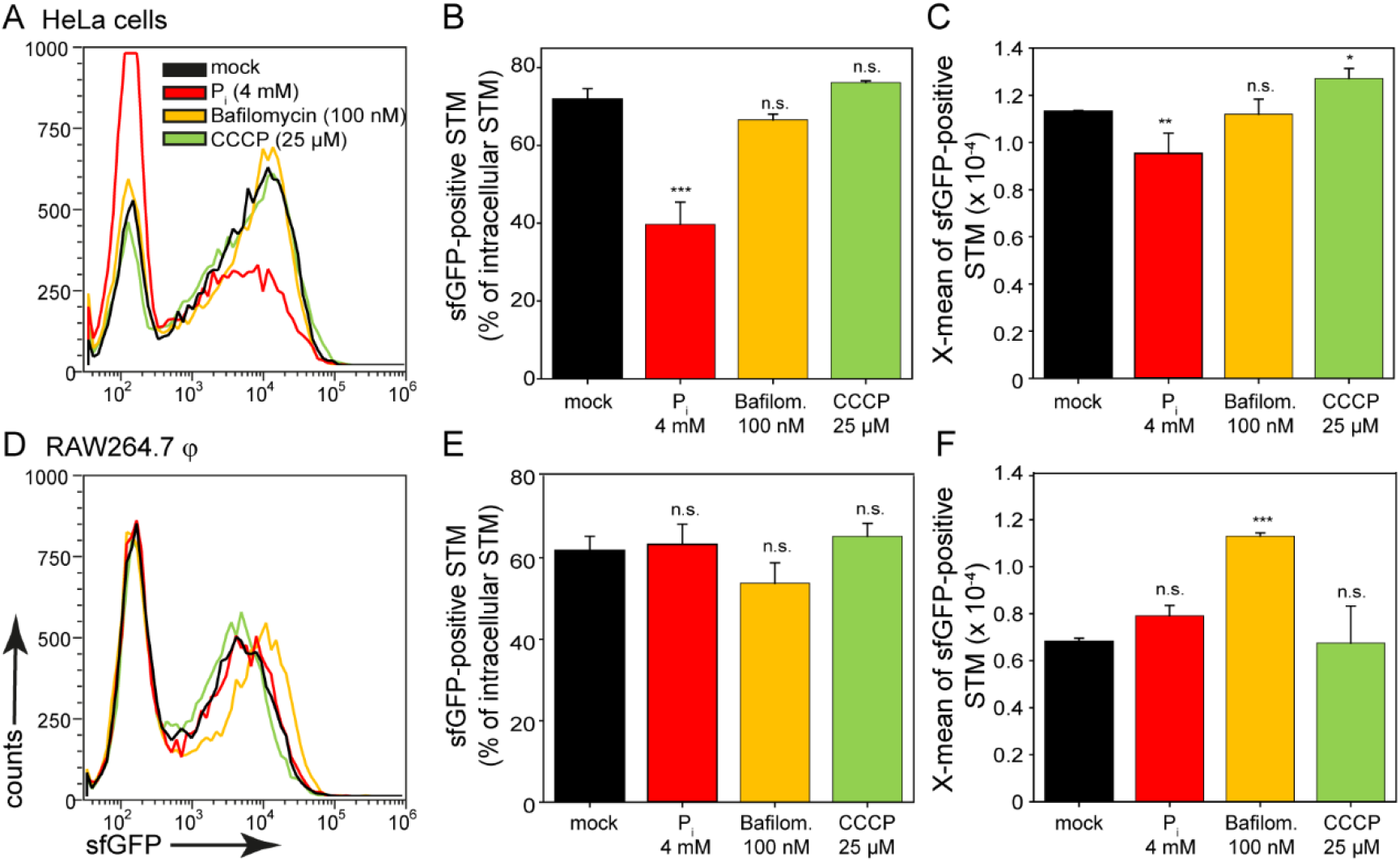
Manipulation of SCV pH and phosphate availability for STM in host cells. HeLa cells (A-C) or RAW264.7 macrophages (D-F) were infected by STM WT [p5440] at MOI 5. Infected cell were left untreated (mock, black), or experimentally manipulated. To alter P_i_ availability, 4 mM P_i_ was added to cells 24 h prior infection by STM and maintained during infection (red). vATPase inhibitor Bafilomycin (orange) or ionophor CCCP (green) were added at 25 μM or 100 nM final concentration, respectively, to infected cells at 1 h p.i., and maintained throughout the infection period. Infected host cells were lysed 8 h p.i., released STM were fixed and subjected to FC for quantification of P_*pstS*_-positive STM (B, E), and the X-means for sfGFP intensities of P_*pstS*_-positive STM (C, F). Means and standard deviations from triplicates are shown. Representative data for STM WT in nontreated and treated HeLa cells (A), or RAW264.7 macrophages (D). Statistical analyses are indicated as for **Figure 3**.

## Discussion

### Main phosphate transporter PstSCAB-PhoU is required for intracellular replication of STM

This study investigated nutrient availability for intracellular STM with focus on phosphate. P_i_ is essential for many important functions like DNA and RNA synthesis and energy metabolism. We developed and applied a sensitive quantitative approach to investigate, at a single cell level, the availability of P_i_ within the intracellular environments of STM. Our study demonstrates the importance of intracellular P_i_ homeostasis for virulence of STM. Deletions of *phoB*, *phoR*, *phoU,* or *pstSCAB* encoding the ABC transporter led to a reduced replication of STM in macrophages and epithelial cells. Several previous studies on intracellular STM reported upregulation of genes such as *pstSCAB* or *phoBR* (Eriksson, Lucchini, Thompson, Rhen, & Hinton, 2003; Garcia-del Portillo et al., 1992; Hautefort et al., 2008). Deletion of *pstSCAB* was recently reported to attenuate systemic virulence of STM in the murine infection model (Zhang et al., 2019), and our work extends these findings to further components of the phosphate metabolism. The reduced replication of a Δ*phoB* strain in RAW264.7 macrophages and a reduced systemic virulence in mice was recently reported (Jiang et al., 2020). Defect of PhoU leads to constitutive expression of the Pst transporter and unphysiological accumulation of P_i_ in STM cytosol, indicating that the proper phosphate homeostasis is of critical importance. Defects in phosphate metabolism also impaired virulence of other pathogens such as *Shigella flexneri*, *Yersinia* spp., or *E. coli,* and several studies have shown that the Pho regulon is part of a complex network important for bacterial virulence and stress response (Lamarche et al., 2008).

Using a dual fluorescence reporter, we measured P_i_ availability and concentrations in distinct intracellular habitats of STM. PstS is a periplasmic protein that binds P_i_ with high affinity and the promoter of *pstS* was repressed in culture by P_i_ concentration above 1 mM, as described for *E. coli* (Rosenberg, Gerdes, & Chegwidden, 1977), while a phosphate down-shock experiment revealed P_i_ concentration above 100 μM as repressing for PstS. Increasing expression of P_*pstS*_∷GFP with decreasing phosphate concentrations was reported for *Shigella flexneri* and in addition a very strong induction measured by concentration of 10 μM P_i_ (Runyen-Janecky & Payne, 2002). Using STM as reporter, we determined P_i_ concentration of less than 10 μM within the SCV of HeLa cells and RAW264.7 macrophages based on the P_*pstS*_∷sfGFP intensity. However, in the cytosol of HeLa cells, concentrations above 10 μM were determined (**Figure 10**A). The released bacteria indicated a heterogeneous distribution and a pronounced change over time. Remarkable was the shift at 10 h p.i. in HeLa cells, where the cytosolic population became smaller in relation to the vacuolar population. At this time, the Δ*ssaV* strain differed from STM WT probably due reduced replication in the SCV (Hensel et al., 1998). In contrast, in RAW264.7 macrophages, strong P_i_ deficiency was measured shortly after invasion. After 10 h p.i., a decreased percentage of P_*pstS*_-positive STM and induction of sfGFP intensity was obvious. We therefore assumed that non-induced STM consisted of metabolically inactive, temporarily cytosolic, as well as dead and dormant bacteria (Helaine et al., 2014; Röder & Hensel, 2020). In activated RAW264.7 macrophages and in primary human macrophages, both exhibiting higher antimicrobial activity (Rosenberger & Finlay, 2002; Schwan, Huang, Hu, & Kopecko, 2000), the induced subpopulation was smaller compared to STM WT in resting RAW264.7 macrophages, and indicated availability of less than 10 μM P_i_.

**Figure 10.**
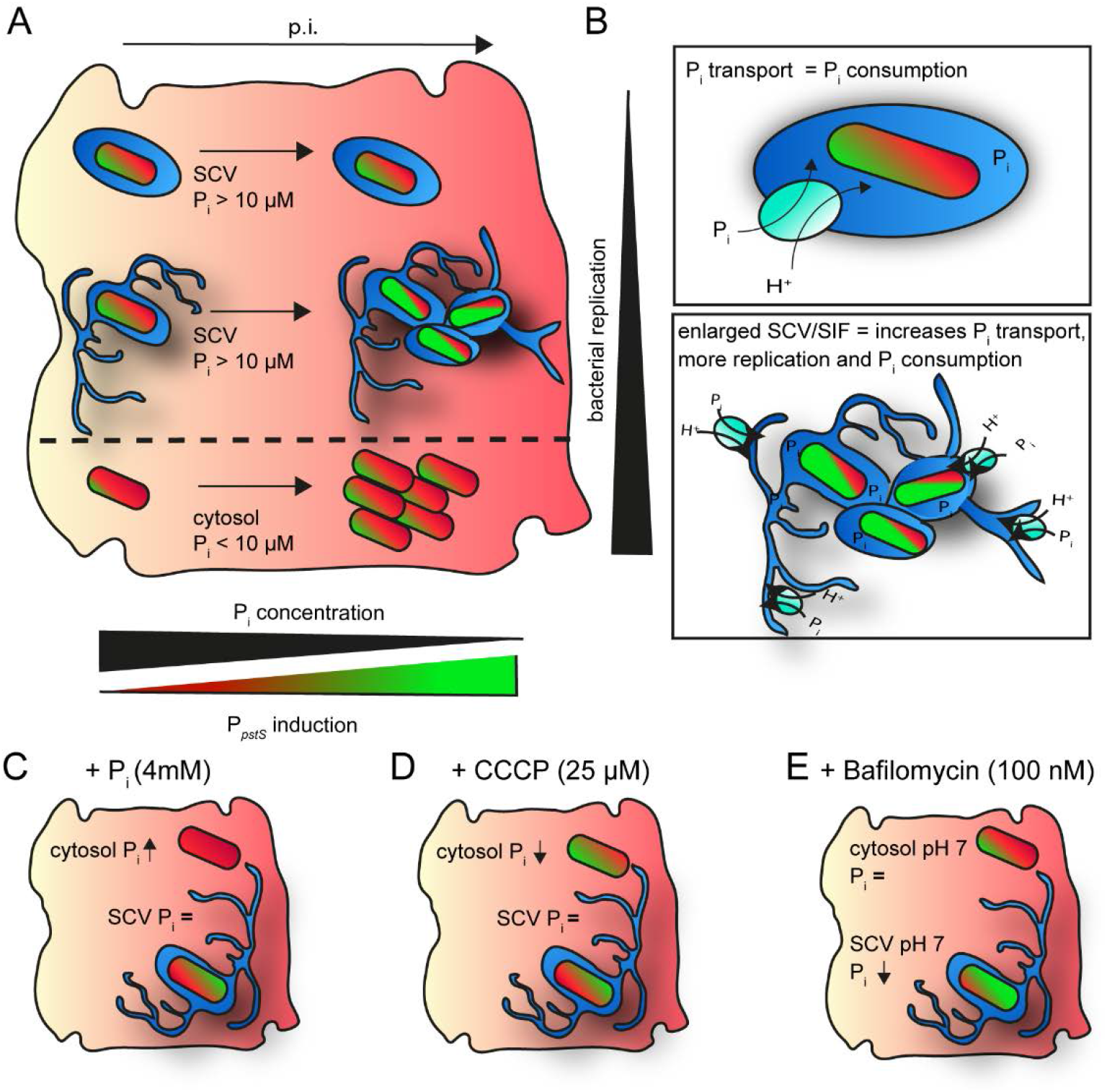
Model of factors that influence P_*pstS*_ induction of STM in host cells. A) The P_*pstS*_ induction depends on the vacuolar or cytosolic presence of STM, and the level of intracellular replication. B) Replication-arrested STM demand less phosphate. An enlargement of the SCV-SIF continuum leads to an increased phosphate import, but also to increased replication and phosphate consumption. C) External addition of phosphate led to increased intracellular phosphate concentration in host cell cytosol, but not within SCV. D) Addition of CCCP led to decreased intracellular phosphate concentration in host cells cytosol, but not within the SCV. Bafilomycin changed the pH within the SCV and led to lower phosphate availability.

Serovars of *S. enterica* cause systemic infections in humans, and macrophages are important for immunity to infection. However, what renders macrophages permissive or restrictive for survival and replication of *S. enterica* serovars is only partially understood. Human M1 macrophages (classically activated) were used here that restrict proliferation of STM (Lathrop et al., 2018). Based on analyses of fluorescence reporter strains in human macrophages, together with the results obtained by treatment with Cotrimoxazole to inhibit the intracellular replication, we conclude that consumption of and demand for P_i_ was lower due to the lack of replication in these habitats, similar to an Δ*ssaV* strain. All these results clearly showed that P_i_ homeostasis allows STM to survive and replicate intracellularly. Therefore, *pstSCAB-phoU* seems to be the main transporter of P_i_ and essential for replication and intracellular survival. Inhibition of such important transporter, or interference with proper sensing and regulation may be considered as new non-antibiotic strategy.

### Multiple effects of phosphate in regulation of virulence factors

It is known that low levels of P_i_ in the SCV trigger the activation of SPI2 genes (Lober, Jackel, Kaiser, & Hensel, 2006). The translocation of effectors via the SPI2-T3SS leads to the maturation of the SCV and the formation of the SIFs (Rajashekar et al., 2008). In RAW264.7 macrophages and HeLa cells, we ascertained a correlation between SPI2 activity and P_*pstS*_ activity. Nearly all bacteria with P_*ssaG*_ induction were also positive for P_*pstS*_ induction (Figure S 4; Figure S 5B). In HeLa cells a further P_*pstS*_-positive population without SPI2 activity was present, mainly the cytosolic subpopulation. Additionally, the PhoBR two-component system represses *hilA* expression under low extracellular phosphate conditions (Lucas et al., 2000). Therefore, low phosphate levels are important to control *hilA* expression in the intestinal environment (Baxter & Jones, 2015). The transcriptional activator HilA controls expression of SPI1-T3SS, as well as some effector proteins for invasion. This may explain reduced invasion of the Δ*phoU* strain. Deletion of *phoU* resulted in a missing activation of the SPI1-T3SS due to the permanent repression of *hilA* by *phoBR*. Therefore, *phoBR* and *pstSCAB-phoU* have two critical roles for virulence, i) to switch off SPI1-T3SS after invasion due to the low P_i_ values in the intracellular space, and ii) to activate SPI2-T3SS, thus enabling the replication of STM during intracellular lifestyle.

Recently, PagR was identified as *S. enterica*-specific integrator of low P_i_ and low Mg^2+^ levels leading to activation of expression of SPI2-T3SS genes (Jiang et al., 2020). The study proposed that PhoB is activated by low P_i_, and PhoQ in response to low Mg^2+^, both activated PagR. PagR activates SlyA, and SlyA induces expression of *ssrAB*. Increased levels of the two-component system SsrAB then result in increased expression of genes encoding the SPI2-T3SS and cognate effector proteins. This cascade explains an amplification loop resulting in additional copies of SsrAB. However, the sensor SsrA also responds directly to physicochemical signals in the SCV in order to activate expression of SPI2-T3SS functions, and phosphate limitation and mildly acidic pH were identified as signals (Deiwick, Nikolaus, Erdogan, & Hensel, 1999; Lober et al., 2006).

Another study observed that MgtC interacts with PhoR leading to activation of the Pho regulon via PhoB and increased P_i_ uptake (Choi et al., 2019). Interestingly, and in contrast to our findings, inactivation of *phoB*, or interference with MgtC-PhoR interaction resulted in decreased P_i_ transport but increased proliferation in macrophages, and higher virulence in a murine model (Choi et al., 2019). These phenomena were explained by dual roles of MgtC in controlling F_0_F_1_ ATPase and intracellular pH (Lee, Pontes, & Groisman, 2013), and the new interaction with PhoR.

Work by Pontes and Groisman (Pontes & Groisman, 2018) suggested a further regulatory cascade. Starvation of STM in the SCV for Mg^2+^ results in ribosome instability, decreased protein biosynthesis, and decreased ATP hydrolysis leading to lower levels of P_i_ in the cytosol of STM. This situation induces the Pst transport system, despite sufficient P_i_ availability in the extracellular space. This model also supports the function of PhoR as sensor for P_i_ in the bacterial cytosol, in line with the absence of the periplasmic domain in PhoR for sensing P_i_ in periplasm and thus extracellular milieu (Gardner & McCleary, 2019).

Future work has to clarify the interconnected effects of extracellular and cytosolic P_i_ concentration, transport, the P_i_ consumption by anabolism during STM proliferation, and the effect of ATP hydrolysis. For this, STM reporter strains are useful tools, yet need to complemented by single cells sensors such as P_i_-sensitive fluorescent proteins that directly sense phosphate levels.

### STM reporter strains as tool for measuring phosphate concentrations

In further experiments, we aimed to induce specific P_i_ changes in host cells and to measure these using STM reporter strains. Our work suggests that STM may serve as simple and sensitive tool for measuring intracellular P_i_ concentrations. In Caco2BBe1 cells external supply of 4 mM P_i_ increases phosphate uptake. The addition of CCCP reduced intracellular P_i_ levels and Bafilomycin showed no effect in CaCo2BBe1 cells on P_i_ uptake (Candeal et al., 2014). We observed similar results using the dual fluorescence reporter after STM infection of HeLa cells, but not of RAW264.7 macrophages. While in HeLa cells a portion of the intracellular population resides in host cell cytosol, in RAW264.7 macrophages nearly all bacteria remain within the SCV. Therefore, we assume that the addition of 4 mM P_i_ or CCCP changes the cytosolic P_i_ concentration of the host cells, but P_i_ levels in the SCV were unaffected. In addition, Bafilomycin only showed detectable changes in RAW264.7 macrophages, maybe as consequence of the cytosolic population in HeLa cells. The cytosol of HeLa cells is neutral with pH 7-8 (Llopis, McCaffery, Miyawaki, Farquhar, & Tsien, 1998), while the SCV acidifies from pH 6 to pH 4-5 which can be inhibited by using Bafilomycin (Rathman et al., 1996). Therefore, the lack of acidification changes the P_i_ availability in the SCV of RAW264.7 macrophages, suggesting a neutral pH decrease the P_i_ influx into the SCV/SIF-continuum.

### Increase of the SCV-SIF continuum enables further transport of P_i_

In vertebrates, P_i_ homeostasis is mediated by sodium-dependent P_i_ transporters that use the inwardly directed electrochemical gradient of Na^+^ ions. In HeLa cells, the major Na^+^-Pi cotransporter is P_i_T1 (Bon et al., 2018). Gradients are established by the Na^+^-K^+^-ATPase to control P_i_ influx (Virkki, Biber, Murer, & Forster, 2007). It is not known if specific P_i_ transporters exist in endosomal membranes that control luminal P_i_. Previous work by Vorwerk *et al*. (2015) investigated the proteome of host cell endomembranes modified by intracellular STM and identified a mitochondrial phosphate carrier protein (MCPC) (Vorwerk, Krieger, Deiwick, Hensel, & Hansmeier, 2015) that catalyzes the transport of P_i_ by H^+^ cotransport into the mitochondrial matrix. Therefore, we assume that P_i_ influx in the SCV/SIF continuum also requires a H^+^ or Na^+^ gradient. Enlargement of membrane surface and lumen of the SCV-SIF continuum may lead to reduction of H^+^ or Na^+^ luminal concentration, thus increase the gradient resulting in increased H^+^ and P_i_ cotransport. Furthermore, an improved fusion with vesicles though the enlarged membrane also possible to increase P_i_ transport (Liss et al., 2017). Therefore, we conclude that STM WT increases P_i_ transport due to the enlargement of SCV-SIF continuum resulting in bacteria replication. An Δ*ssaV* strain, without the SCV-SIF continuum, cannot replicate due to the lack of access to P_i_ (**Figure 10**B).

In conclusion, we used STM as reporter to monitor, at single cell level, phosphate availability in distinct intracellular niches and host cell types. Proper P_i_ homeostasis is critical for the intracellular lifestyle of STM and the non-redundant nature of the high affinity P_i_ transporter may indicate a new target for therapeutic interference with systemic *S. enterica* infections.

## Acknowledgements

This work was supported by the Deutsche Forschungsgemeinschaft by grant HE 1964/18-2 and BMBF by grant 031L0093A as part of the Infect-ERA cluster SalHostTrop. We thank Jörg Deiwick for construction of p3776, Pascal Felgner for cloning p5081, Monika Nietschke for excellent technical support and Tatjana Reuter for support by the isolation of primary human macrophages.

## Data availability statement

The data that support the findings of this study are available from the corresponding author upon reasonable request.

## Authors contributions

JR and MH conceived the study, JR performed experimental work, JR and MH analyzed the data, JR and MH wrote the manuscript.

## Materials and Methods

### Bacterial strains and growth conditions

*Salmonella enterica* serovar Typhimurium (STM) strains NCTC 12023 (identical to ATCC 14028) and isogenic mutant strains are summarized in Table 1. STM strains were routinely cultured in Luria-Bertani (LB) broth containing 50 μg x ml^−1^ carbenicillin (Roth) if required for selection of plasmids. Bacterial cultures were routinely grown in glass test tubes at 37 °C with aeration in a roller drum at ca. 60 rpm. For invasion of HeLa cells, fresh LB medium was inoculated 1:31 with o/n cultures of STM and incubated for 3.5 h with agitation in a roller drum. To test induction of reporters by inorganic phosphate (Pi), PCN media was supplemented with various amounts of P_i_ ranging from 100 nM to 25 mM.

**Table 1.**
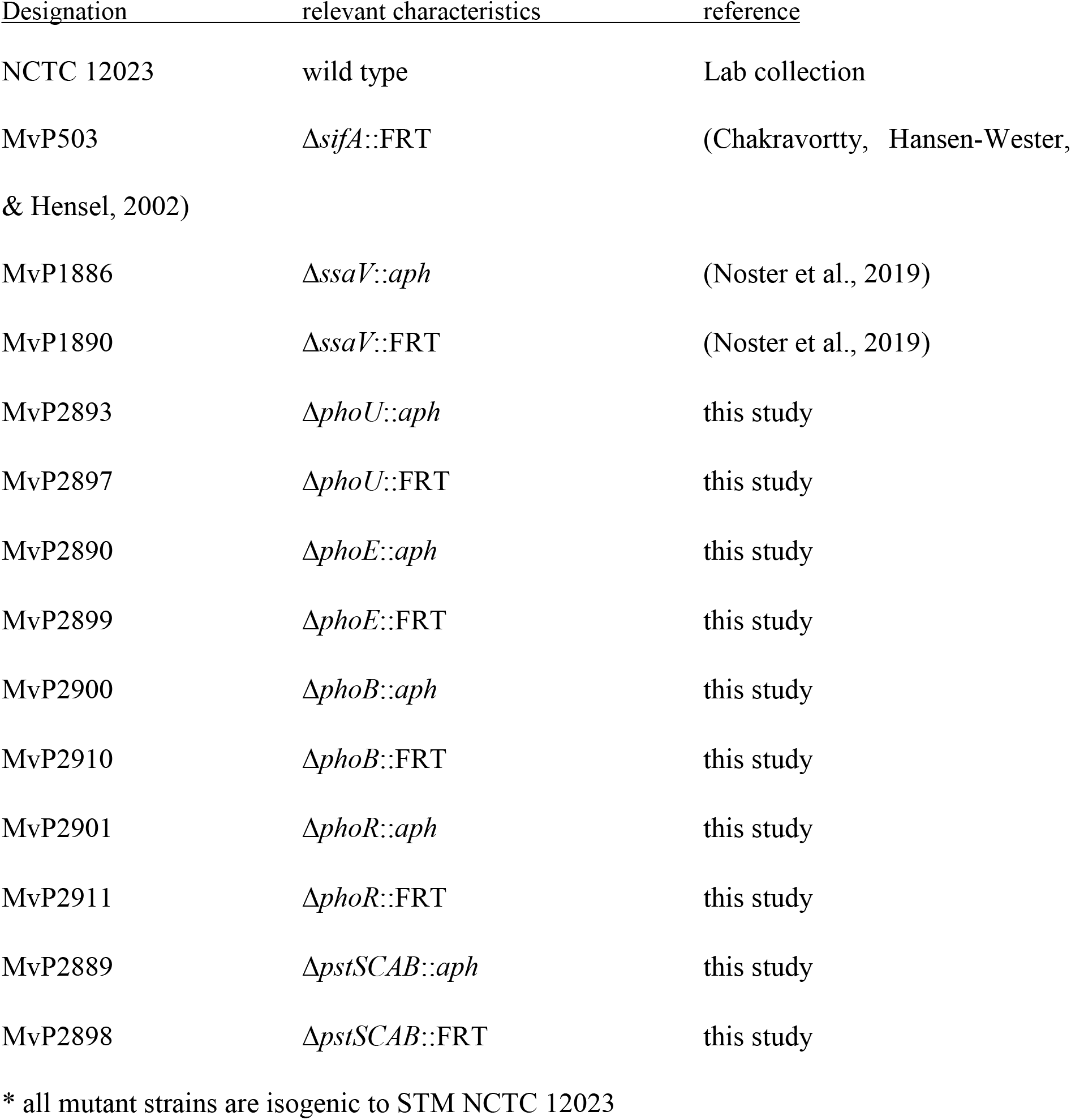
*Salmonella enterica* serovar Typhimurium strains used in this study

### Construction of plasmids

Plasmids used in this work are listed in Table 2. Oligonucleotides for generation of recombinant DNA molecules were obtained from IDT and are specified in Table S1.

**Table 2.**
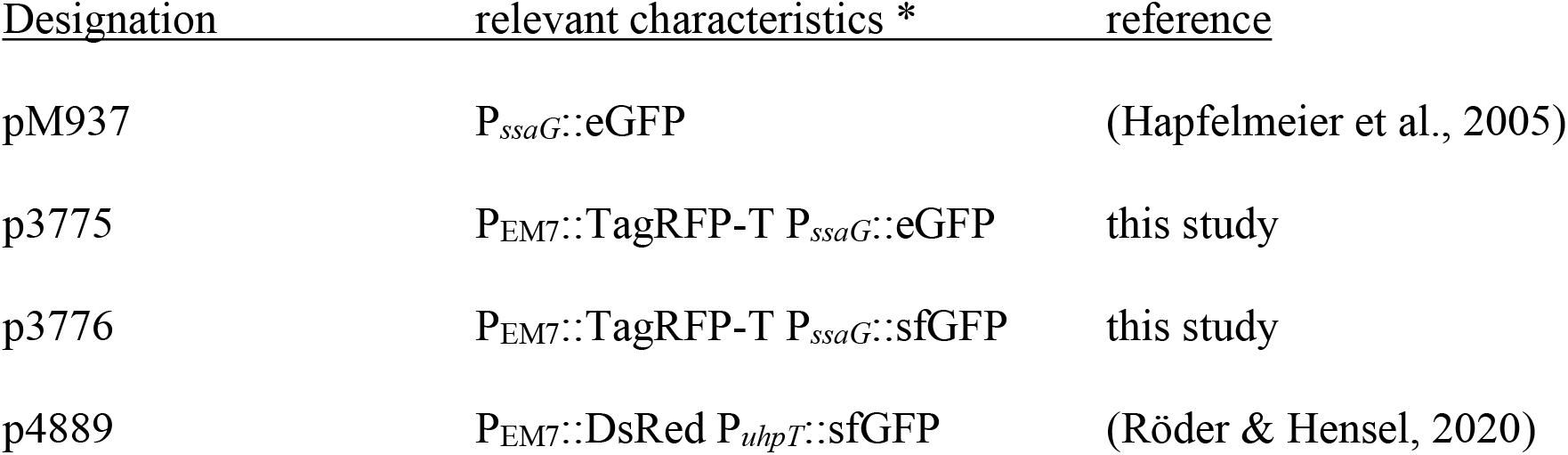
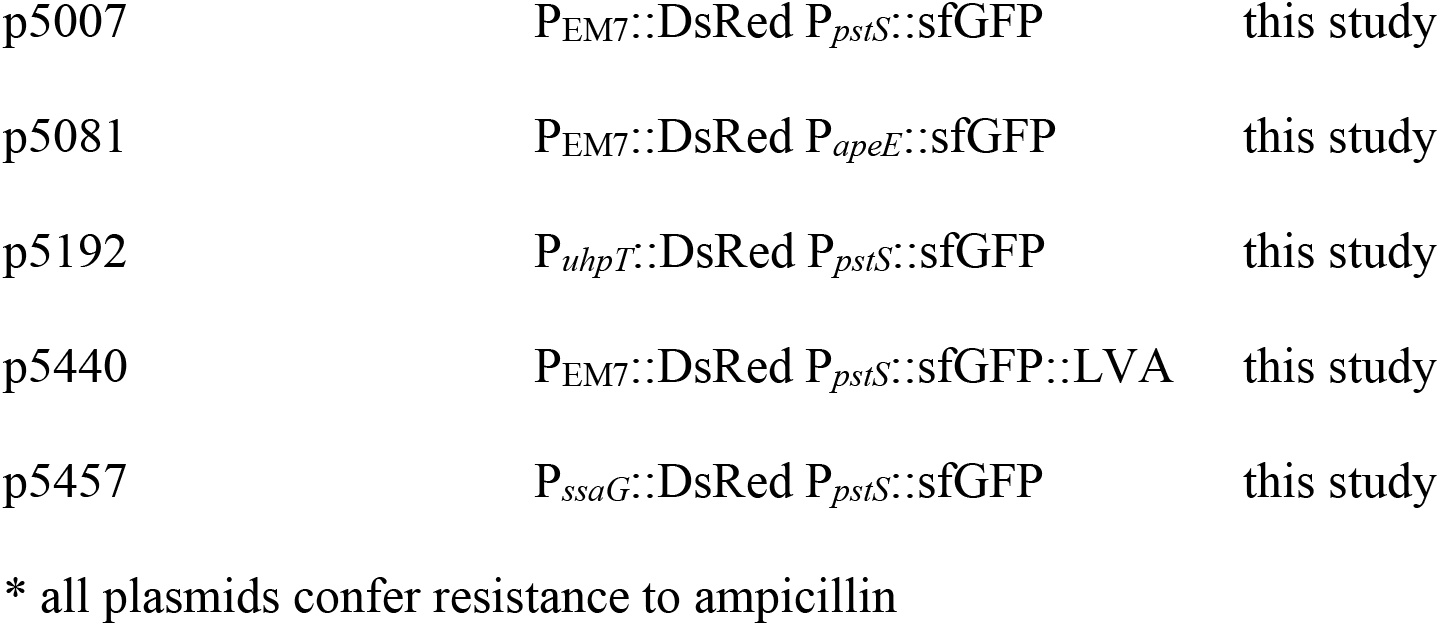
Plasmids used in this study

The promoter of *pstS* or *apeE* of STM was cloned as 300 bp fragment upstream of the translational start site of *pstS* or *apeE* using 1f-P_*pstS*_ and 1r-P_*pstS*_ or 1f-_*apeE*_-p4889 and 1r-_*apeE*_-p4889 for amplification of the insert. Primers Vf-p4889 and Vr-p4889 were used for amplification of vector p4889, and fragments were fused by Gibson assembly (GA) to generate p5007 or p5081. Dual fluorescence reporter p5007 and p5081 harbor P_EM7_∷DsRed for constitutive expression of DsRed, and P_*pstS*_∷sfGFP or P_*apeE*_∷sfGFP for sfGFP expression.

Plasmid pM937 was used to generate p3775 by insertion of transcriptional terminators and P_EM7_∷Tag-RFP-T amplified from pWRG338 using primers P_EM7_-RFP For and RFP-T2 Rev and cloning as NotI/SacI fragment. The eGFP gene in p3775 was replaced by sfGFP obtained as XhoI/BamHI fragment from pWRG167 to generate p3776.

The promoter of *ssaG* was cloned as 320 bp fragment upstream of the translational start site of *ssaG*. EM7 was replaced by P_*ssaG*_ using GA to create p5457 with primers Vf-p4889-exEM7 and Vr-p4889-exEM7 for amplification of vector p5007, and 1f-PssaG-p4889 and 1r-PssaG-p4889 for amplification of the insert from p3776. The promoter of *uhpT* was cloned as 251 bp fragment upstream of the translational start site of *uhpT*. The module P_EM7_∷DsRed was replaced by module P_*uhpT*_∷DsRed using GA to create p5192 with primers Vf-p4889 ex EM7 and Vr-p4889 ex EM7 for amplification of vector p5007, and 1f p5407 PuhpT and 1r-p4507 Dsred for amplification of the insert from p5407.

For destabilization of sfGFP, the LVA tag according to BioBricks (AGGCCTGCTGCAAACGACGAAAACTACGCTTTAGTAGCT) was fused to the C-terminus to sfGFP via SDM with primers p4889-LVA SDM For and sfGFP-LVA Rev to create p5440.

### Cell lines and cultivation

Human epithelial cell line HeLa were maintained in DMEM containing 4.5 g x l^−1^ glucose, 4 mM L-glutamine and sodium pyruvate (Biochrom) supplemented with 10% FCS in an atmosphere of 5% CO_2_ and 90% humidity at 37 °C. The murine macrophage cell line RAW264.7 (ATCC no. TIB-71) were cultured in DMEM containing 4.5 g x l^−1^ glucose and 4 mM stable glutamine (Biochrom) supplemented with 6% FCS.

### Preparation of primary human macrophages

Primary human macrophages were prepared from buffy coat from pooled blood samples of anonymous donors (obtained from the Deutsches Rotes Kreuz) as described in Bonifacino *et al*. Page 2/6 – 2/9 (Bonifacino, Dasso, B., Lippincott-Schwartz, & Yamada, 2004). Preparation of lymphocytes by Ficoll-Hypaque gradient was performed as described, alternatively to whole blood, buffy coat was mixed 1+1 with PBS. For differentiation into monocytes/macrophages, the isolated lymphocytes were thawed, seeded and maintained in RPMI-1640 (Biochrom), supplemented with 20% FCS and 2.5 ng x ml^−1^ GM-CSF (Peprotech). After 5-7 days, the purity of the monocyte/macrophage population was checked by staining with FITC anti-human CD14 antibody (BioLegend) and FC, and subsequently used for infection.

### Infection of host cells

Host cell infections were performed as previously described (Rajashekar et al., 2008). Briefly, RAW246.7 macrophages and human macrophages were infected with o/n cultures, and HeLa cells were infected with 3.5 h subcultures of STM at multiplicity of infection (MOI) of 5 or 25. Infected cells were centrifuged at 500 x g for 5 min to synchronize infection, incubated for 25 min at 37 °C in an atmosphere of 5% CO_2_, before extracellular bacteria were removed by washing thrice with PBS. Subsequently, host cells were maintained in growth media containing 100 μg x ml^−1^ gentamicin for 1h, followed by media containing 10 μg x ml^−1^ for remaining time of incubation.

### CI assay

Competitive Index assay was performed as previously described (Segura, Casadesus, & Ramos-Morales, 2004). Briefly, HeLa cells or RAW264.7 macrophages were seeded in surface-treated 24-well plates (Faust) and grown to 80% confluency at the day of infection. WT without antibiotic resistance and mutant strains harboring the *aph* cassette were separately grown in LB or LB containing 50 μg x ml^−1^ kanamycin. Cells were infected with a 1+1 mixture of WT and mutant strain at an overall MOI of 1 for 25 min. Non-invaded bacteria were killed using media containing 100 μg x ml^−1^ gentamicin for 1 h and replaced by media containing 10 μg x ml^−1^ gentamicin for further incubation. The numbers of viable intracellular bacteria were determined 1 h and 16 h p.i. by plating onto LB agar with and without kanamycin (AppliChem). The CI for bacterial survival is defined as the ratio of x-fold replication of WT to mutant strain for 1 h.

### Pulse-chase with fluid phase markers

For tracing the endocytic pathway, fluid phase markers were used. HeLa cells and RAW264.7 macrophages were incubated with 100 μg x ml^−1^ AlexaFluor 647-conjugated dextran, (molecular weight 10,000, Molecular Probes) o/n, washed, and incubated with dextran-free media. Cells were infected for 25 min, incubated 1 h with 100 μg x ml^−1^ gentamicin, cultivated 15 h in growth media with a decreased gentamicin concentration of 10 μg x ml^−1^ and processed for imaging.

### Live cell imaging

HeLa cells and RAW267.4 macrophages cultured in 8-well chamber slides were infected at MOI of 5. The infection was performed as described above and 16 h p.i., the medium was replaced by Imaging Medium supplemented with 10 μg x ml^−1^ gentamicin. Live cell imaging was carried out by CLSM on a Leica SP5 with an environmental incubation chamber maintaining a humidified atmosphere of 5% CO_2_.

### Quantification by flow cytometry analyses

HeLa cells and RAW264.7 macrophages were infected at MOI of 5 for 25 min. At various time points from 2 to 24 h p.i., cells were lysed with 0.1% Triton X-100 and fixed with 3% PFA for subsequent FC analyses using the Attune NxT Cytometer (Thermo Fischer). Experiments were performed in triplicates at least three times. Data were analyzed with Attune NxT 2.5. Statistical analyses were performed using One-Way Anova Bonferroni using SigmaPlot 13 (Systat Software).

## Suppl. Materials Captions

**Figure S 1:**
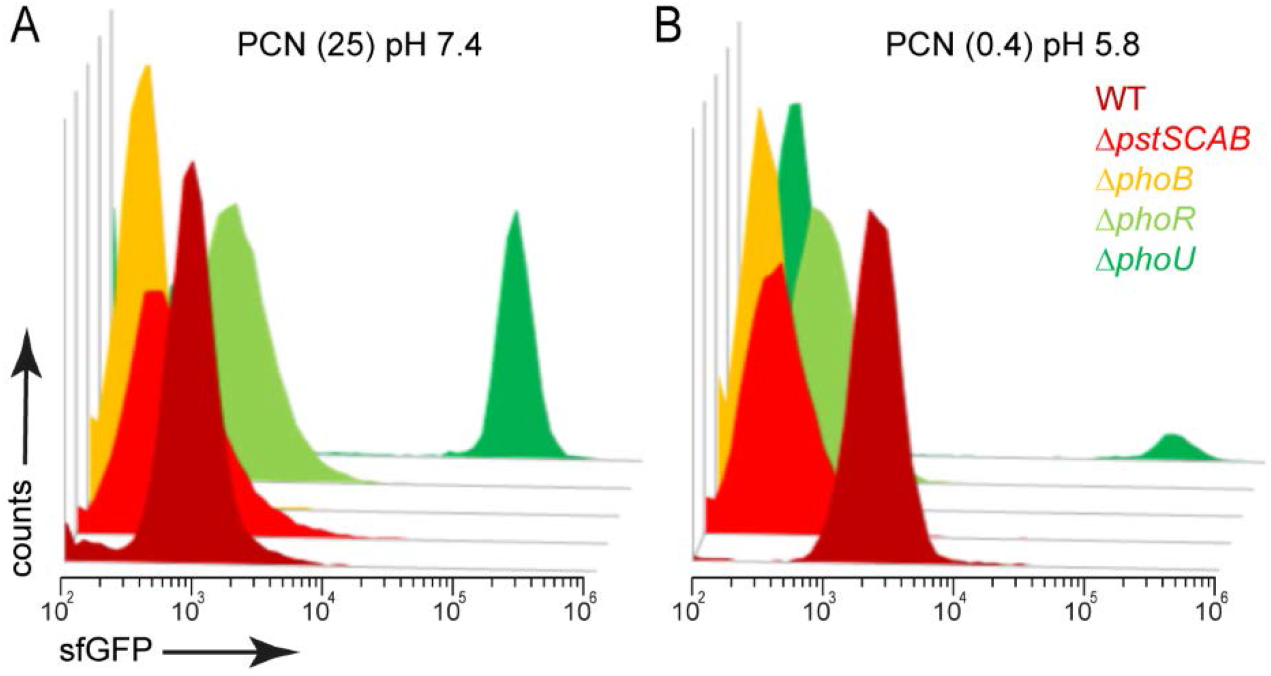
The phosphate reporter P_*pstS*_∷sfGFP is responsive to defects in PstSCAB, PhoB, PhoR, or PhoU. STM WT, Δ*pstSCAB*, Δ*phoB*, Δ*phoR, or* Δ*phoU* strains, all harboring p5007 were grown in PCN minimal medium (A) at pH 7.4 containing 25 mM P_i_ (PCN (25) pH7.4), or (B) at pH 5.8 containing 0.4 mM P_i_ (PCN (0.4) pH 5.8). Samples were collected after 3.5 h of culture and sfGFP intensity of P_*pstS*_-induced bacteria was determined by flow cytometry (FC).

**Figure S 2:**
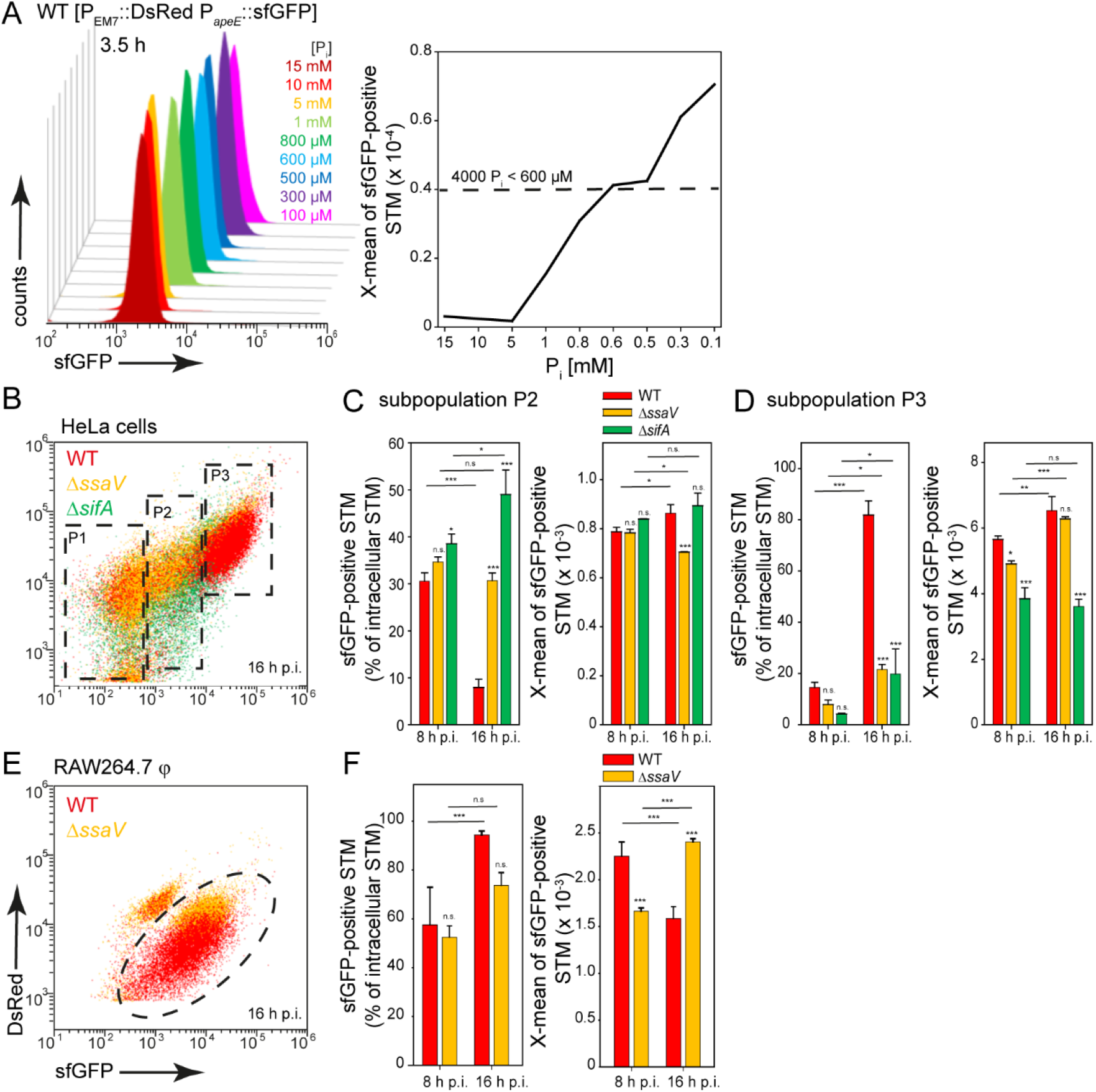
Performance of an alternative dual fluorescence phosphate reporter based on P_*apeE*_∷sfGFP. STM strains harbored p5081 for constitutive expression of DsRed and sfGFP under control of P_*apeE*_. A) Induction of P_*apeE*_∷sfGFP *in vitro* during growth in presence of various phosphate concentrations [P_i_]. STM WT [p5007] was grown o/n in PCN (25), pH 7.4 and then diluted 1:31 in PCN, pH 7.4 with various concentrations of P_i_ as indicated. Samples were collected after 3.5 h of subculture and sfGFP intensities of P_*apeE*_-induced bacteria was determined by FC. HeLa cells (B-D) or RAW264.7 macrophages (E, F) were infected at MOI of 5 with STM WT (red), Δ*ssaV* (orange) or Δ*sifA* (green) strains as indicated, each containing the phosphate reporter p5081. The host cells were lysed 8 h or 16 h p.i., and released STM were fixed. Subsequently, the bacteria were subjected to FC to quantify the size of the induced intracellular population and X-means of sfGFP intensity of the P_*apeE*_-induced bacteria. The bacterial population was divided into 3 subpopulations based on P_*apeE*_ intensity. Representative data for STM WT, Δ*ssaV* and Δ*sifA* strains at 8 h and 16 h p.i. in HeLa cells (C, D), and for STM WT and Δ*ssaV* strains at 8 h and 16 h p.i. in RAW264.7 macrophages (F). Mean values and standard deviations of the P_*apeE*_-positive bacteria populations from triplicates of a representative experiment are shown. Statistical analyses are indicated as for **Figure 3**.

**Figure S 3:**
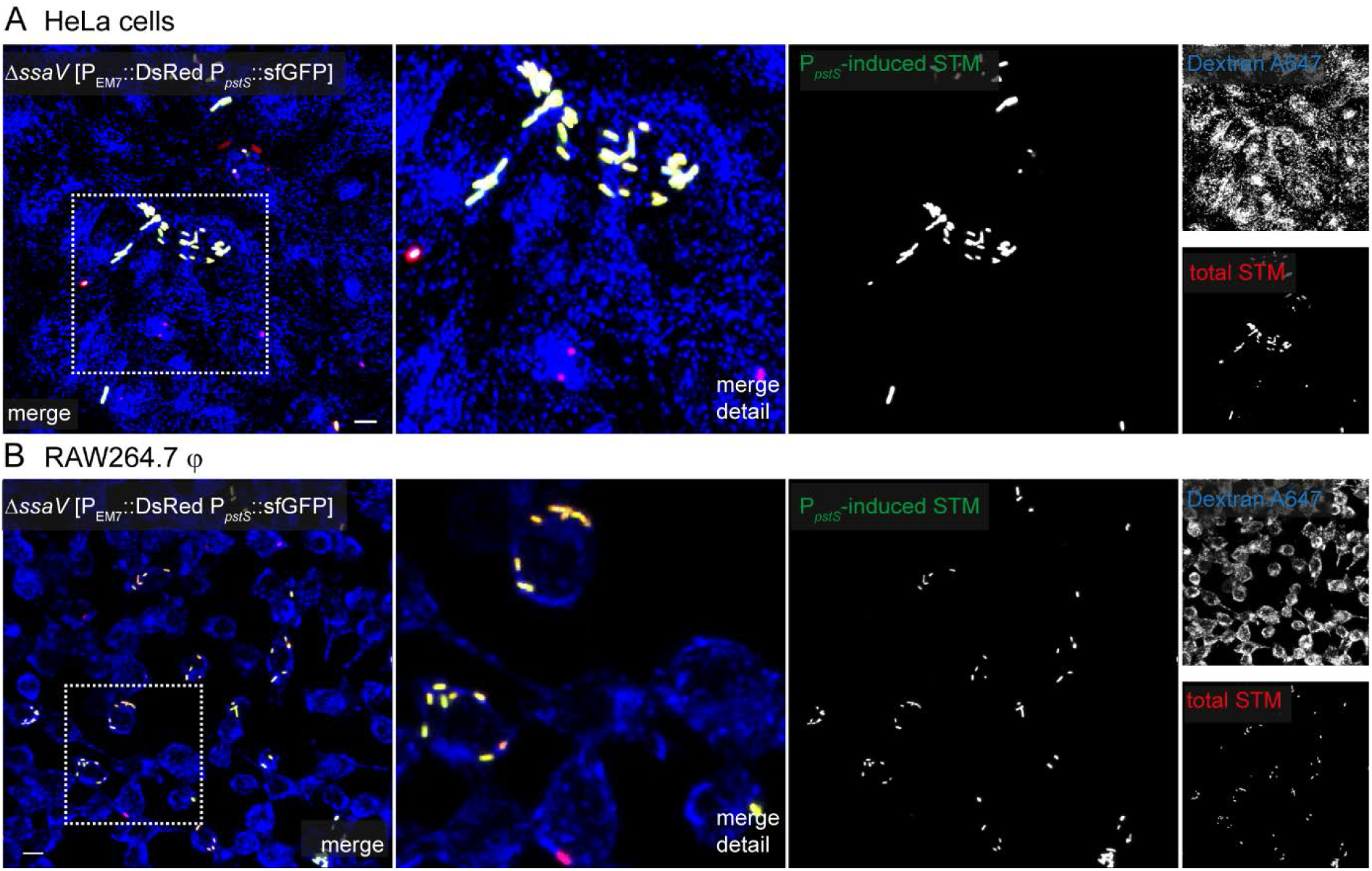
Heterogeneous P_*pstS*_-induced sfGFP intensity of intracellular STM Δ*ssaV*. HeLa cells (A), or RAW264.7 macrophages (B) were infected by STM Δ*ssaV* [p5007] and pulse-chased with dextran-AlexaFluor 647 for labelling of the endosomal system. Live cell imaging was performed 16 h p.i. Representative infected cells indicate red and green fluorescence signal for STM. Scale bars, 10 μm.

**Figure S 4:**
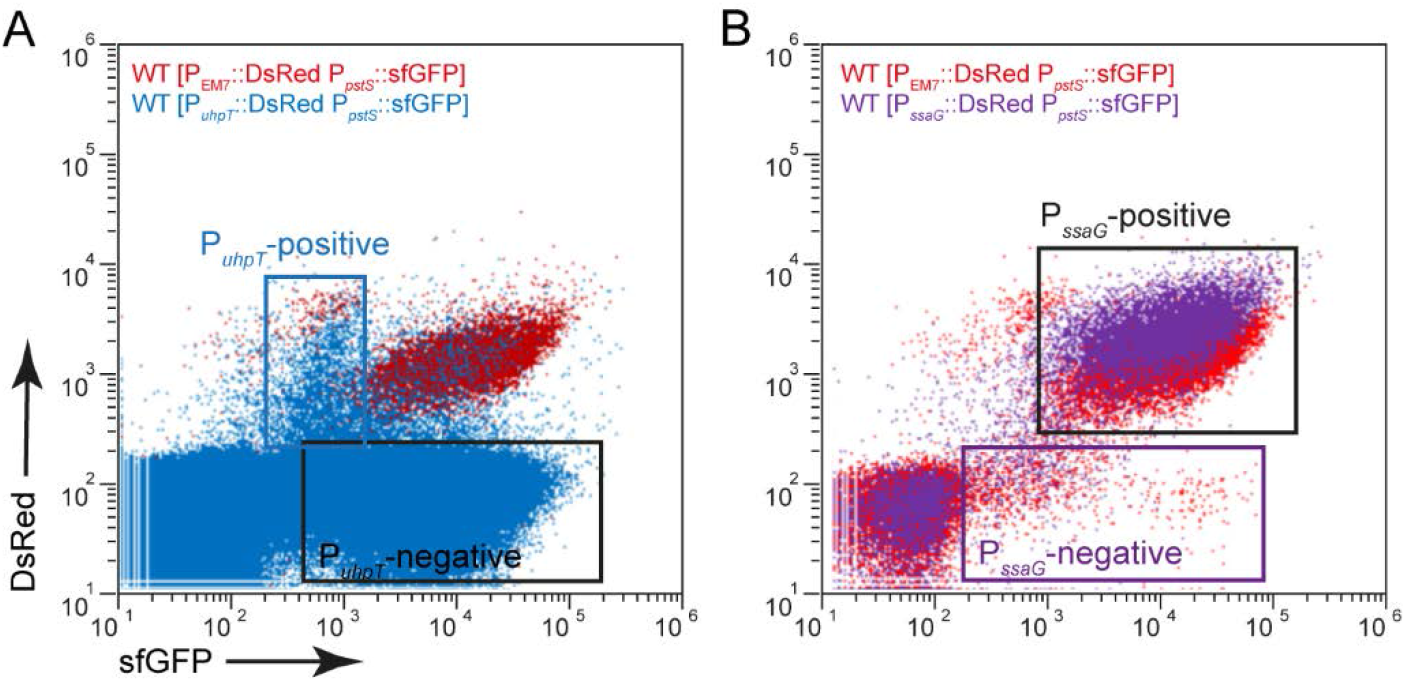
Cytosolic bacteria are rarely found in RAW264.7 macrophages. RAW264.7 macrophages were infected with STM WT [p5193] (blue) or STM WT [p5457] (violet) at a MOI of 5. For comparison and gating of populations, cells were infected with STM WT [p5007] (red). At 16 h p.i., host cells were lysed, released bacteria fixed and subjected to FC for quantification of intracellular STM positive for P_*uhpT*_, P_*ssaG*_, and/or P_*pstS*_. Representative data are shown for (A) STM WT [p5193], and (B) STM WT [p5457].

**Figure S 5:**
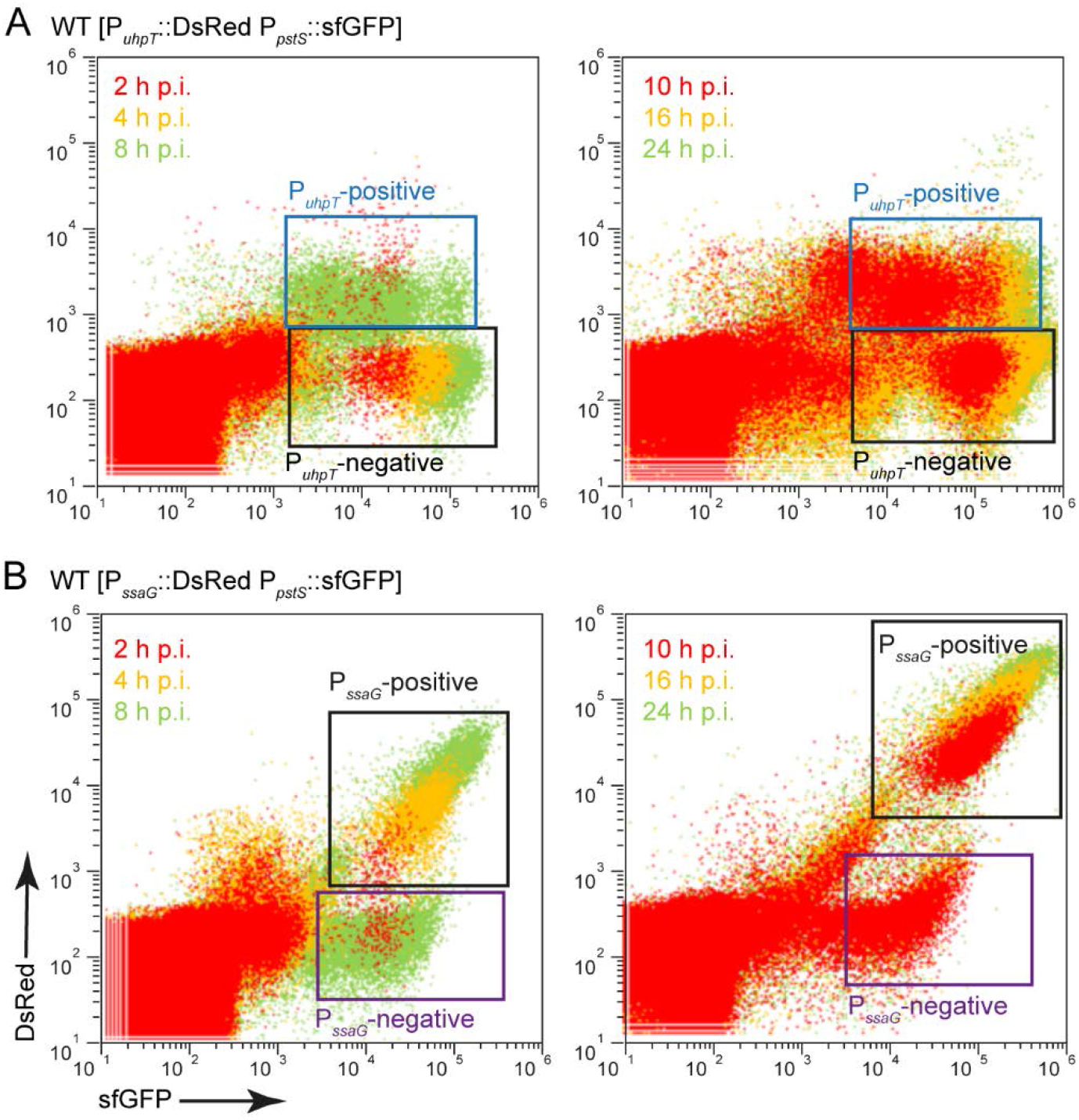
Cytosolic STM encounter higher phosphate concentration than SCV-bound bacteria. HeLa cells were infected with (A) STM WT [p5193], or (B) STM WT [p5457] at a MOI of 5. At various time points p.i., host cells were lysed, released bacteria fixed and subjected to FC for quantification of P_*pstS*_-positive bacteria in combination with P_*uhpT*_ or P_*ssaG*_ induction. Representative data for (A) STM WT [p5193] indicate the P_*uhpT*_∷sfGFP-positive cytosolic population, and (B) STM WT [p5457] indicate the P_*ssaG*_∷sfGFP-positive SCV-bound population.

**Figure S 6:**
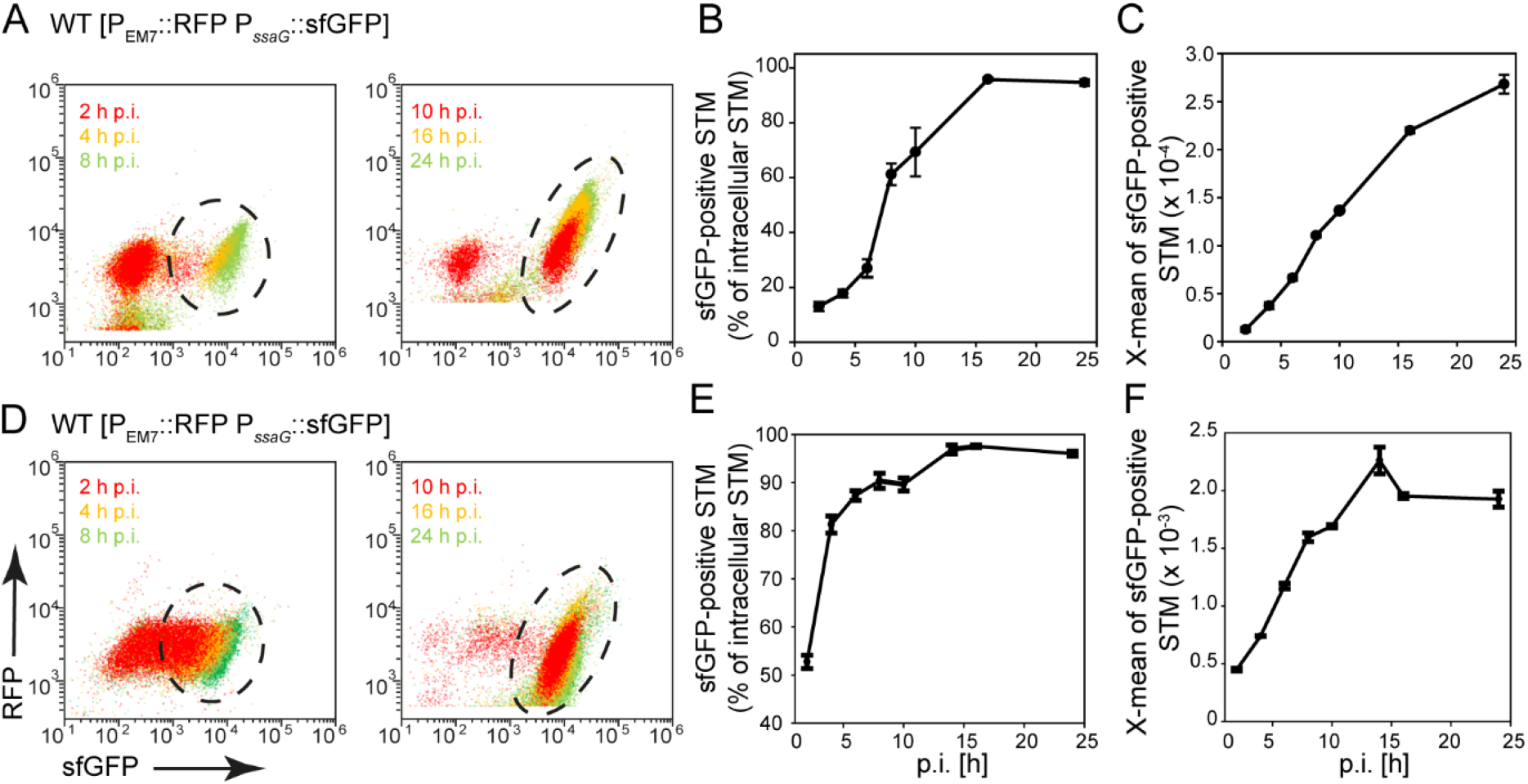
SPI2-T3SS activity increases in HeLa cells and RAW macrophages. STM WT harboring p3776 for constitutive expression of RFP and sfGFP under control of P_*ssaG*_ was used to infect HeLa cells (A-C) or RAW264.7 macrophages (D-F) at MOI of 5. At various time points p.i., host cells were lysed, released STM were fixed and subjected to FC for quantification of P_*ssaG*_-positive bacteria and sfGFP intensity. Means and standard deviations for the size of the P_*ssaG*_∷sfGFP-induced population (B, E) and X-means of sfGFP-fluorescence (C, from triplicates are shown for various time points p.i. Representative data for STM WT in HeLa cells (A) or in RAW264.7 macrophages (D).

**Figure S 7:**
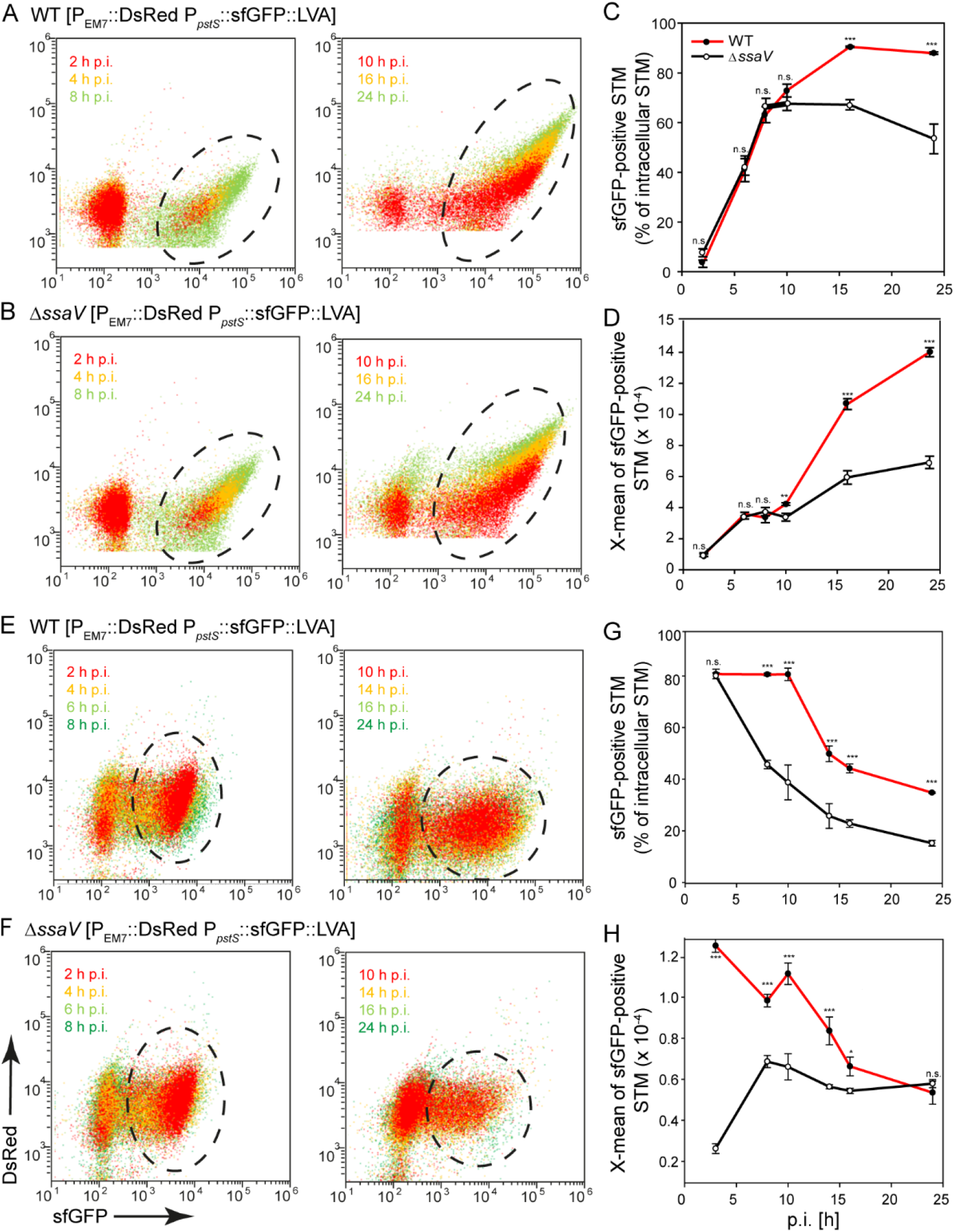
Time-resolved changes in phosphate availability in HeLa cells and RAW264.7 macrophages. HeLa cells or RAW264.7 macrophages were infected with STM WT (A, C, E, G, red lines) or Δ*ssaV* (B, D, F, H, black lines) strains harboring phosphate reporter p5440 at a MOI of 5. Host cells were lysed at various time points p.i., released STM were fixed and subjected to FC to quantify P_*pstS*_-positive bacteria and the sfGFP intensity over time in HeLa cells or RAW264.7 macrophages. Mean values and standard deviations indicate the size of the sfGFP-positive populations (C, G) and the X-means of sfGFP intensities (D, H) of triplicates at various time points p.i. as indicated. Representative data for STM WT and Δ*ssaV* strains in HeLa cells (A, B) or RAW264.7 macrophages (E, F) are show for various time points p.i. Statistical analyses are indicated as for **Figure 3**.

**Table S1.**
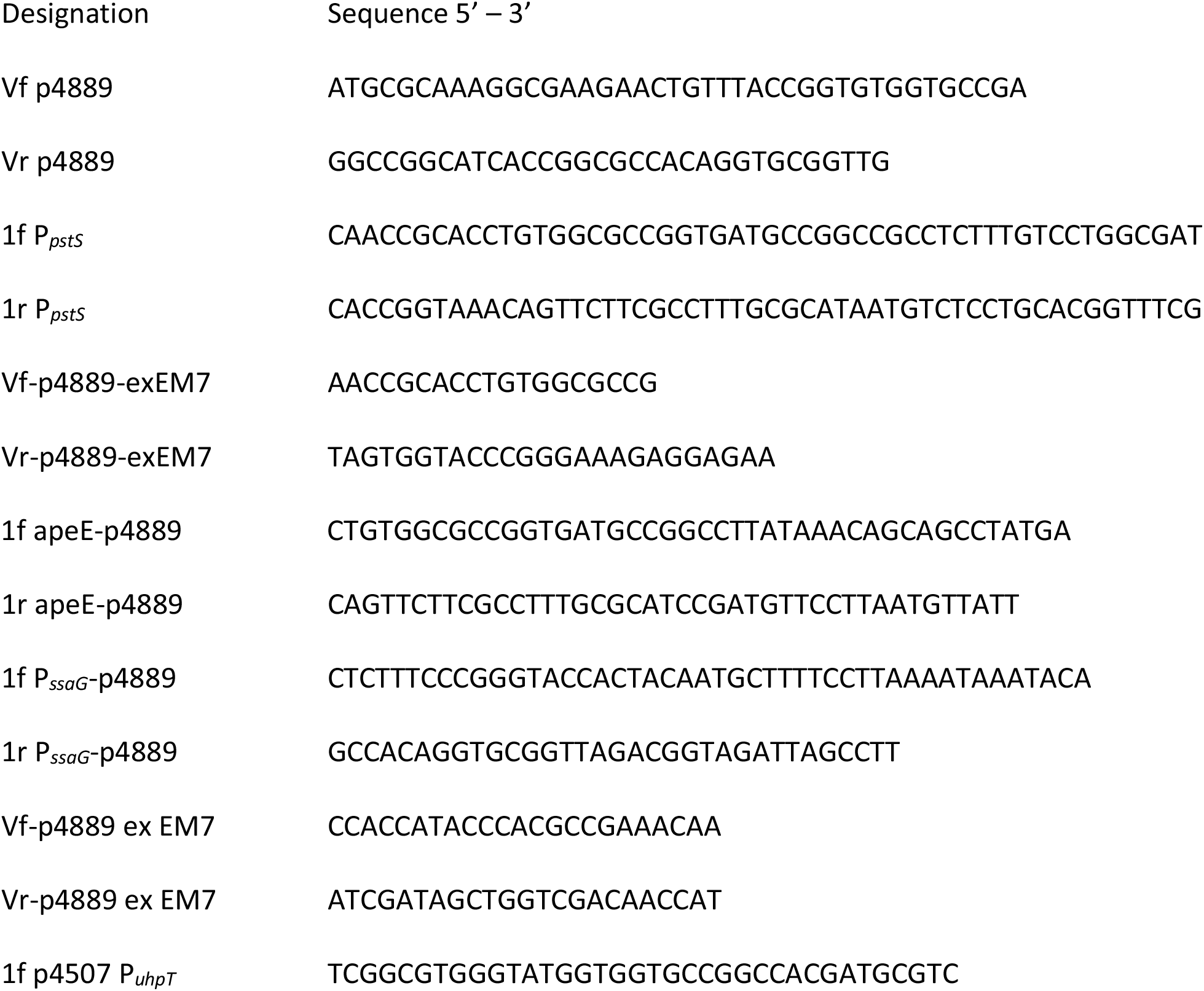
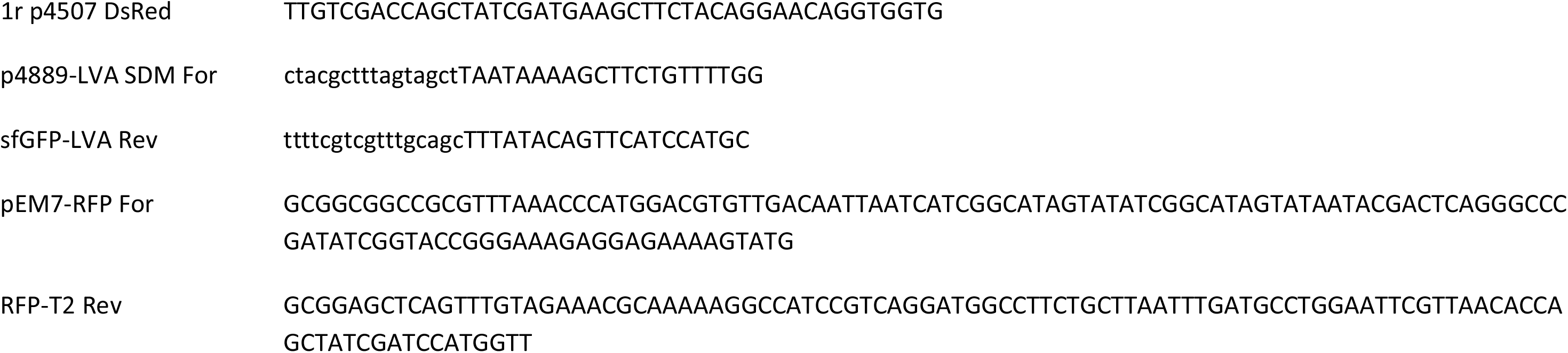
Oligonucleotides used in this study

